# Chelator-Free Radiometal Labeling Inside Engineered Affibodies

**DOI:** 10.64898/2026.01.28.702152

**Authors:** Lani J. Davies, Frank Bruchertseifer, Alfred Morgenstern, Sarah Spreckelmeyer, Christoph Nitsche

## Abstract

Affibodies are remarkably stable three-helix bundle proteins that can be engineered to selectively bind target proteins. When combined with radioactive metals, they serve as imaging agents or cancer therapeutics, depending on the metal used. Traditionally, this involves bifunctional linkers that attach large chelators to the affibody via reactive groups. Here, we present an alternative approach that eliminates the need for such linkers by burying the metal within the core of the affibody, surrounded by its three helices. A simple engineered triple cysteine motif, with one cysteine in each helix, stably binds Bi(III), Pb(II), In(III) and Ga(III), which are commonly used in imaging and radiotherapy. Quantitative metal uptake is instantaneous at room temperature and physiological pH, and all metal-affibody complexes remain fully intact for one week at 4 °C. All retain their metal cargo when challenged with cellular concentrations of glutathione, while only the bismuth-affibody complex withstands a challenge with 100 equivalents of strong chelators, even over two weeks. We demonstrate selective uptake and retention of ^213^Bi, a promising isotope for targeted alpha therapy.

## Introduction

Affibodies are small, engineered proteins based on a three-helix bundle structure derived from the Z domain, a stabilized variant of the antigen-binding B domain of *Staphylococcal* protein A (Figure 1A).^[1-4]^ Developed through directed evolution, they combine extraordinary stability under extreme pH, temperature and chemical denaturation with strong binding affinity and high specificity to their targets. Their compact size (6–7 kDa, 58 amino acids) allows excellent tissue penetration, making affibodies valuable tools for molecular imaging and targeted cancer therapy, where the short half-life of radioisotopes and systemic circulation of targeting agents must be matched.^[5]^ Unlike immune-derived antibodies, affibodies can be readily produced via recombinant microbial expression or chemical synthesis, enabling site-specific incorporation of unnatural amino acids, fluorophores or chelators as well as direct fusion to the solvent-exposed termini.^[3, 5]^ In molecular imaging, affibodies have emerged as tracers for HER2-specific imaging as promising alternative to antibodies and nanobodies (single-domain antibodies).^[6]^ Unlike full-length antibodies, which face challenges with short-lived positron emitters due to their long systemic circulation, affibodies can generate clear images of tumors within one hour after administration and provide high tumor-to-background contrast.^[7-8]^

**Figure 1.**
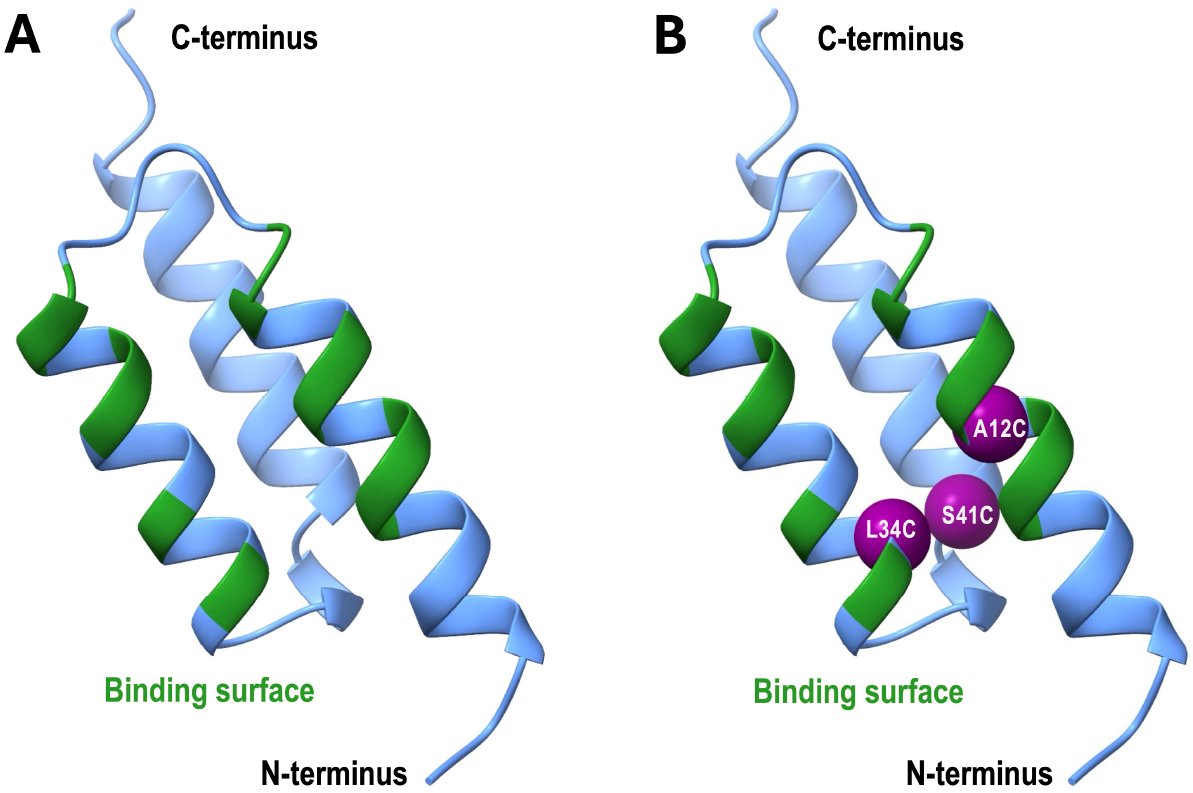
Metal‐binding affibody design. (A) Backbone ribbon structure of the wildtype affibody Z_HER2:2891_ with residues involved in target binding highlighted in green. (B) Backbone ribbon structure of the engineered affibody Z_HER2:2891_‐3C showing mutations A12C, L34C and S41C (purple spheres), forming a buried triple‐cysteine motif for radiometal labeling.

The conventional approach for attaching radionuclides to targeting proteins such as antibodies, nanobodies or affibodies involves the use of bifunctional linkers.^[9-10]^ These linkers contain on one side a reactive group designed to bind to cysteine or lysine side chains of the biomolecule and on the other side a chelator binding the metal of interest. This strategy presents significant challenges, particularly in achieving selective tagging and efficient and stable chelator loading with the radiometal, which often requires harsh conditions such as elevated temperatures.^[10-12]^ In addition, the chelator itself can affect the properties of the targeting protein. The large and highly charged nature of many chelators can substantially alter the overall physicochemical characteristics of the radiolabeled biomolecule. This effect is especially pronounced in small proteins like affibodies, where the chelator significantly affects size and charge.

Affibodies have been conjugated to bifunctional linkers for theranostic development and have progressed to clinical trials.^[13]^ For example, ^111^In-or ^68^Ga-labeled DOTA^0^-Z_HER2:342-pep2_ (ABY-002) can produce high-quality images of tumors as early as two hours post-injection.^[14]^ An engineered variant with higher thermal stability (ABY-025), also conjugated to ^111^In via the same 1,4,7,10-tetraazacyclododecane-1,4,7,10-tetraacetic acid (DOTA) chelator, has likewise shown promise as a HER2-specific labeling agent.^[15]^ However, synthesis is complex and upscaling remains challenging, limiting broader clinical application. By removing the need for bulky bifunctional chelators, we demonstrate direct metal binding to the affibody structure and further improve their already excellent properties.

## Results and Discussion

Previously, we engineered nanobodies to bind diagnostic and therapeutic metals by exploiting their natural disulfide bond and introducing cysteine mutations within 4–8 Å of it, creating triple- and quadruple-cysteine binding motifs.^[16]^ Building on this concept, we designed a triple-cysteine motif within the three helices of an affibody. Because affibodies are naturally cysteine-free (Figure 1A), we used an AlphaFold2-predicted structure to identify one non-antigen-binding residue in each helix for cysteine substitution (Figure 1B).^[17-18]^ Our earlier work showed that buried multiple-cysteine motifs can bind Bi(III), In(III) and Ga(III) after partial protein denaturation and refolding, covering key applications in targeted alpha therapy (TAT), single-photon emission computed tomography (SPECT) and positron emission tomography (PET), respectively.^[16]^ Here, we aim to further extend this to Pb(II), as ^212^Pb is a β-emitter and an *in vivo* generator of the short-lived α-emitter ^212^Bi.^[19]^ For the first time, we also explore the loading and stability of the radioactive isotope ^213^Bi, an α-emitter used in TAT.^[20]^

We investigated two therapeutically relevant affibodies for this study: Z_HER2:2891_, which targets the HER2 receptor overexpressed in breast and other cancers,^[21]^ and Z_TNF‐alpha1_, which binds the inflammatory cytokine TNFα.^[22]^ Both wildtype constructs and their corresponding triple‐cysteine mutants (Z_HER2:2891_‐3C, Z_TNF‐alpha1_‐3C) expressed in *E. coli* in high yield (Figure 2G,H) and demonstrated helical folding in circular dichroism (CD) spectroscopy (Figure 2C,D). Assuming that partial denaturation would be required to fully reduce the cysteine residues buried within the helix bundle,^[16]^ we initially reduced the affibodies at acidic pH with excess tris(2-carboxyethyl)phosphine (TCEP) for up to 60 minutes at a temperature 8–10 °C below their thermal denaturation midpoint, before subsequently exploring milder conditions (Figure 2I). The optimal and most gentle conditions for near-quantitative labeling of both affibodies with all four metals were 25 °C, pH 7.5 with excess TCEP, as confirmed by native mass spectrometry (Figure 2A,B,I). This streamlined metal‐uptake protocol is well‐suited to clinical workflows, enabling efficient preparation of affibodies without harsh pretreatment. Furthermore, native mass spectrometry (MS) of the wildtype affibodies treated under these optimized conditions confirmed that metal binding is specific to the engineered triple‐cysteine motif and not due to nonspecific interactions with the His_6_-affinity tag (Figure 2J,K). Folding and denaturation midpoints remain largely unchanged following metal uptake (Figure 2C-F). NMR experiments with uniformly ^15^N-labeled Z_HER2:2891_‐3C show only minor peak perturbations upon addition of Bi(III), confirming that the overall affibody fold is preserved (Figure S41).

**Figure 2.**
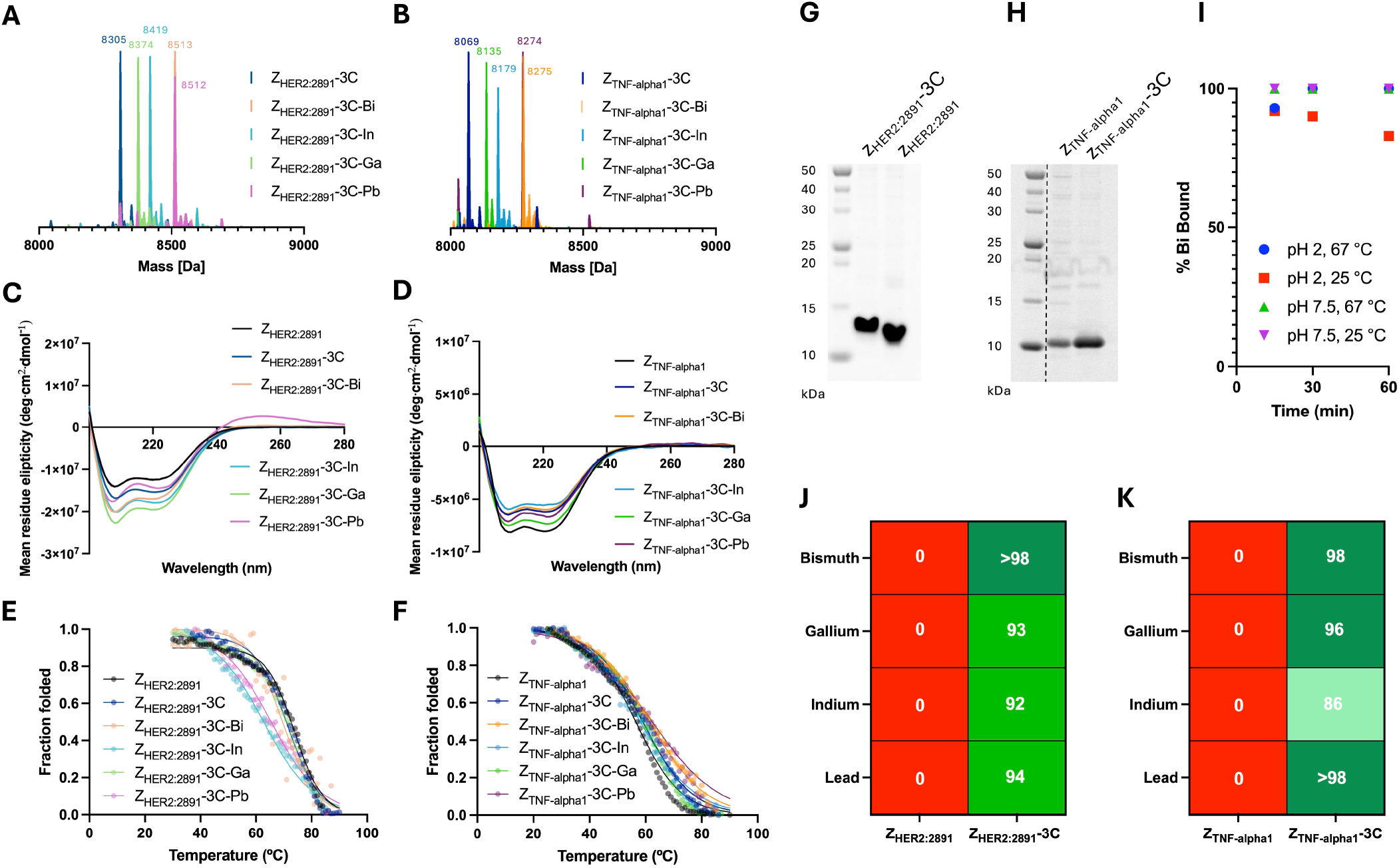
Reduction and metal uptake of engineered affibodies. (A, B) Superimposed native mass spectra (MS) of Z_HER2:2891_-3C and Z_TNF-alpha1_-3C obtained before and after modification with Bi(III), In(III), Ga(III) or Pb(II) in ammonium acetate buffer, pH 7.0. Affibodies were reduced with 50 mM TCEP in 10 mM Tris-HCl, pH 7.5, 150 mM NaCl for 60 min at 25 °C prior to addition of 5 equivalents of either gastrodenol (bismuth tripotassium dicitrate), InCl_3_, Ga(NO_3_)_3_ or Pb(NO_3_)_2_. (C, D) Circular dichroism (CD) spectra in phosphate buffer, pH 7.5 of Z_HER2:2891_-3C (C) and Z_TNF-alpha1_-3C (D) before and after modification with Bi(III), In(III), Ga(III) or Pb(II) compared to their wildtypes. (E, F) Thermal denaturation curves determined by CD spectroscopy in phosphate buffer, pH 7.5 of Z_HER2:2891_ (T_m_ = 74 °C), Z_HER2:2891_-3C (T_m_ = 71 °C), Z_HER2:2891_-3C–Bi (T_m_ = 69 °C), Z_HER2:2891_-3C–In (T_m_ = 63 °C), Z_HER2:2891_-3C–Ga (T_m_ = 71 °C), Z_HER2:2891_-3C–Pb (T_m_ = 65 °C), Z_TNF-alpha1_ (T_m_ = 56 °C), Z_TNF-alpha1_-3C (T_m_ = 59 °C), Z_TNF-alpha1_-3C–Bi (T_m_ = 61 °C), Z_TNF-alpha1_-3C–In (T_m_ = 58 °C), Z_TNF-alpha1_-3C–Ga (T_m_ = 59 °C) and Z_TNF-alpha1_-3C–Pb (T_m_ = 62 °C). (G, H) SDS-PAGE of Z_HER2:2891_ (G) and Z_TNF-alpha1_ (H) constructs after recombinant expression from *E. coli* and His_6_-tag affinity purification. (I) Time-dependent reduction and modification of Z_HER2:2891_-3C with 50 mM TCEP and 5 equivalents of Bi(III) from gastrodenol at different indicated pH and temperatures. Bi(III) uptake (%) was determined by native MS. (J, K) Uptake (%) of Bi(III), Ga(III), In(III) and Pb(II) by affibodies containing no (Z_HER2:2891_ and Z_TNF-alpha1_) or three (Z_HER2:2891_-3C and Z_TNF-alpha1_-3C) cysteine residues after reduction with 50 mM TCEP, pH 7.5 for 60 min at 25 °C and subsequent addition of 5 equivalents of respective metal salts, determined by native MS in ammonium acetate buffer, pH 7.0.

Next, we examined the stability of the metal-labeled affibodies. Z_HER2:2891_‐3C loaded with Bi(III), In(III), Ga(III) or Pb(II) remained fully intact for at least seven days at 4 °C (Figure 3A). To probe complex stability under more challenging conditions, we exposed the Z_HER2:2891_‐3C–metal conjugates to large excesses of physiological and synthetic metal binders, including glutathione (GSH; Figure 3B) and ethylenediaminetetraacetic acid (EDTA; Figure 3C). GSH, the most abundant intracellular thiol, had little effect with all four conjugates remaining 80–90% saturated after 1 hour at 5 mM GSH. The EDTA challenge produced clearer differences. Bismuth binding was only marginally affected at 100 equivalents (∼90% saturation), whereas indium, gallium and lead saturation were significantly reduced (Figure 3C). After 1 hour with 10 equivalents of EDTA, indium and lead were fully removed and only ∼25% of gallium remained bound. Because analogous peptide-Bi(III) complexes do not withstand this level of EDTA challenge,^[23-24]^ these results indicate exceptional kinetic stability of the bismuth-affibody complex.^[25]^

**Figure 3.**
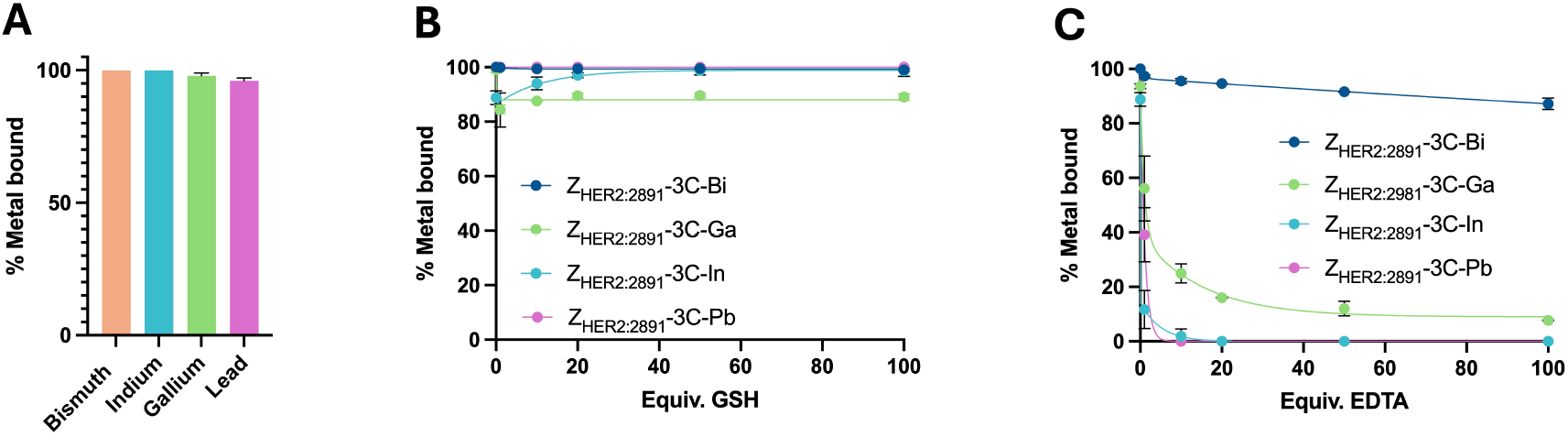
Stability of metal-affibodies determined by native MS. (A) Retention of Bi(III), In(III), Ga(III) and Pb(II) in Z_HER2:2891_-3C after 7 days of storage at 4 °C in ammonium acetate buffer, pH 7.0 (*n* = 3, ± SD). (B, C) Retention of Bi(III), In(III), Ga(III) and Pb(II) bound to Z_HER2:2891_-3C as a function of increasing equivalents of GSH (B) and EDTA (C) after 1 h incubation at 25 °C in ammonium acetate buffer, pH 7.0 (*n* = 2, ± SD).

To further investigate the exceptional stability of the bismuth-affibody complex, we examined it under even harsher conditions (Figure 4). Z_HER2:2891_‐3C–Bi remained ∼80% intact even in presence of 100 equivalents of the stronger chelator diethylenetriaminepentaacetic acid (DTPA; Figure 4A). Investigations of the kinetic stability showed that the Z_HER2:2891_‐3C–Bi complex remained largely intact even after two weeks in a 100‐fold excess of GSH and EDTA (Figure 4B). Orthogonal mass spectrometry further confirmed the stability of Z_HER2:2891_‐3C–Bi (Figure S39) and repeating the EDTA competition with Z_TNF‐alpha1_‐3C–Bi yielded the same high retention of bismuth (Figure S34). Even heating Z_HER2:2891_‐3C–Bi to 100 °C for 10 minutes did not cause any metal release (Figure 4C).

**Figure 4.**
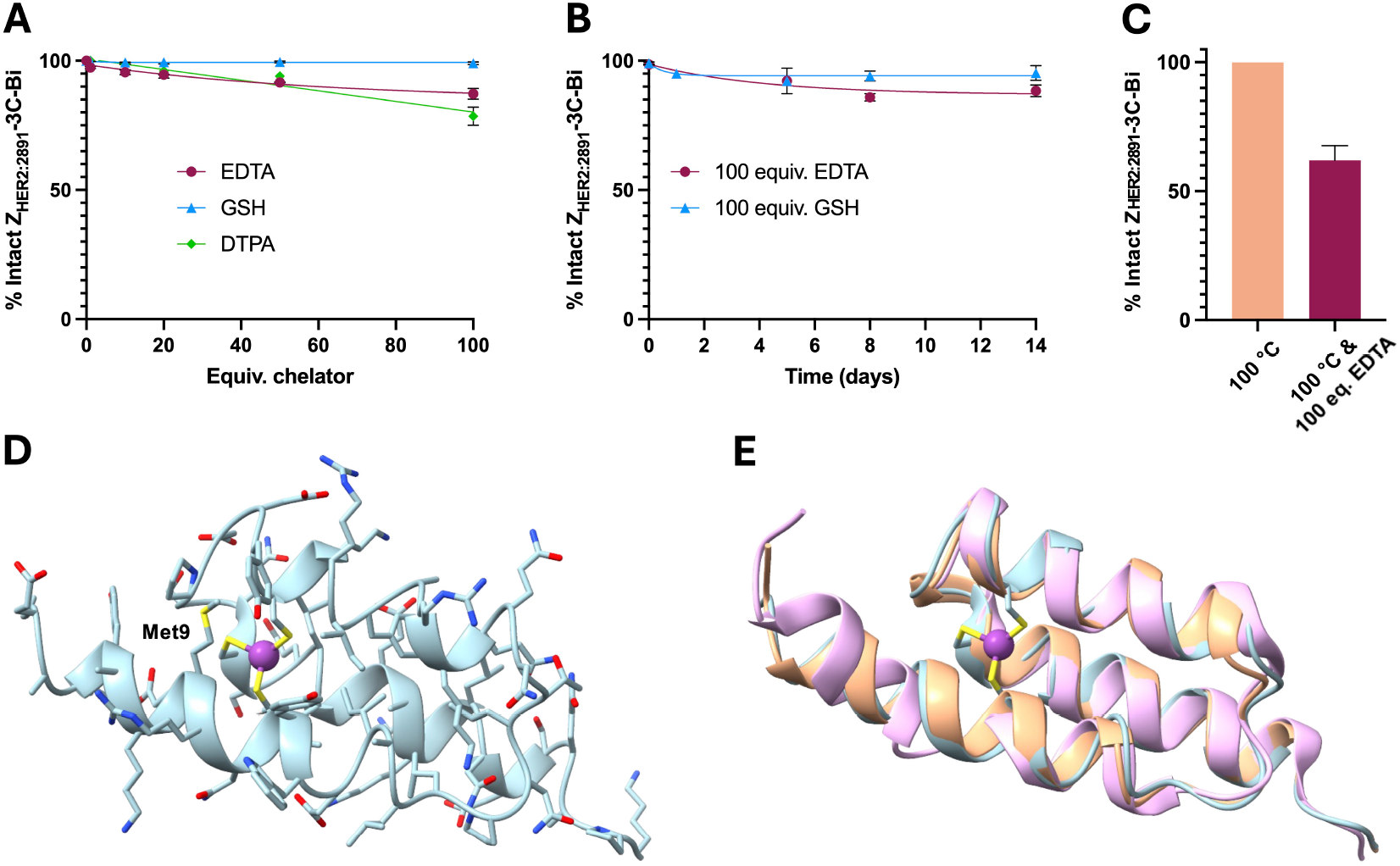
Stability and proposed structure of bismuth-affibodies. (A,B) Retention of Bi(III) bound to Z_HER2:2891_-3C in the presence of increasing equivalents of GSH, EDTA or DTPA after 1 h of incubation at 25 °C (A), and as a function of increasing incubation time in the presence of 100 equivalents of GSH or EDTA at 4 °C (B), in ammonium acetate buffer, pH 7.0 (*n* = 2, ± SD) determined by native MS. (C) Retention of Bi(III) bound to Z_HER2:2891_-3C after heating at 100 °C for 10 min in the absence (orange) and presence (maroon; *n* = 2, ± SD) of 100 equivalents of EDTA, as determined by native MS. (D) Computational model of Z_HER2:2891_-3C–Bi with three cysteine residues (yellow) coordinated to one Bi(III) atom (purple) in a trigonal pyramidal geometry. Side chains are colored by heteroatom type (oxygen, red; nitrogen, blue; sulfur, yellow). (E) Superimposed backbone (ribbon representation) of Z_HER2:2891_-3C–Bi (blue; bismuth, purple; sulfur, yellow), an AlphaFold2-generated structure of wildtype Z_HER2:2891_ (orange) and the solution NMR structure of parent affibody Z_HER2:342_ (pink; PDB code: 2ZKI).

While all four metal-affibody complexes were stable, the exceptional behavior of the Bi(III) conjugate became apparent only under strong chelator challenge. In(III) and Ga(III) are harder Lewis acids and therefore more effectively competed by the hard Lewis base EDTA, whereas Bi(III) is better matched to the soft thiolate donors of the engineered triple-cysteine site.^[26]^ The fact that In(III) and Ga(III) form negatively charged complexes with cysteine-based ligands potentially imposes additional constraints on the coordination geometry.^[27]^ Although Pb(II) and Bi(III) are isoelectronic and soft and both support trigonal‐pyramidal thiolate coordination, Bi(III) forms a neutral Bi(SR)_3_ complex, whereas Pb(II) forms the anionic [Pb(SR)_3_]^−^ species with cysteines,^[28]^ providing a potential explanation for the observed differences in chelator resistance, although this strong difference remains unexpected.

The high retention of Bi(III) in our engineered affibodies under challenge with 100 equivalents of EDTA or DTPA prompted further investigation. A three‐dimensional model of Z_HER2:2891_‐3C–Bi, fully supporting the affibody fold, suggests that the Bi(III) ion is buried within the affibody core, limiting access by external chelators (Figure 4D,E). Additional coordination from methionine-9 (Figure 4D) may further stabilize the complex and help explain the greater resistance of Bi(III) relative to the other metals studied. Consistent with our buried-site hypothesis, thermal denaturation of Z_HER2:2891_‐3C–Bi at 100 °C for 10 minutes in the presence of 100 equivalents of EDTA resulted in partial metal loss (∼60% saturation), indicating that unfolding exposes Bi(III) to the chelator (Figure 4C).

The extreme complex stability of the Bi(III)-affibody conjugates under chelator challenge and heat suggests suitability for application as therapeutic radiopharmaceutical, where stable radionuclide retention is essential. In addition, the clear preference of Bi(III) for the soft thiolate environment of the affibody, together with the weaker binding of harder metals such as Ga(III) and In(III), indicates a level of selectivity that conventional chelators cannot provide. This raises the possibility that mother and daughter isotopes with differing metal hardness could be separated during the complexation step, something that is almost impossible with routinely used chelators, which typically show little discrimination between metals. In addition, it is desirable to separate expression and purification of the affibody from the metal‐loading step, since bismuth isotopes such as ^213^Bi have very short half‐lives (46 minutes). We therefore explored whether the engineered affibody could be supplied as a lyophilized powder that can be stored and shipped at room temperature, then reconstituted with water and charged with the radioactive metal directly in the preclinical setting.

We lyophilized Z_HER2:2891_‐3C in 10 mM Tris-HCl, pH 7.5, containing 150 mM NaCl and stored the resulting powder at room temperature. Reconstitution with water restored the original affibody and buffer concentration. Subsequent reduction with TCEP and Bi(III) loading using the standard protocol yielded fully bismuth‐charged Z_HER2:2891_‐3C–Bi, which remained soluble and correctly folded, as confirmed by native MS, CD spectroscopy and thermal denaturation analysis (Figure 5A–C). LC–MS analysis under acidic conditions (0.1% formic acid) and significant acetonitrile content still showed a single peak corresponding to the intact bismuth‐affibody complex, with full metal retention (Figure 5D). Notably, synthetic affibodies purified by similar HPLC conditions (0.1% TFA) have similarly been reported to retain their native fold and bioactivity,^[29]^ consistent with our observation that the Bi(III) complex withstands these conditions.

**Figure 5.**
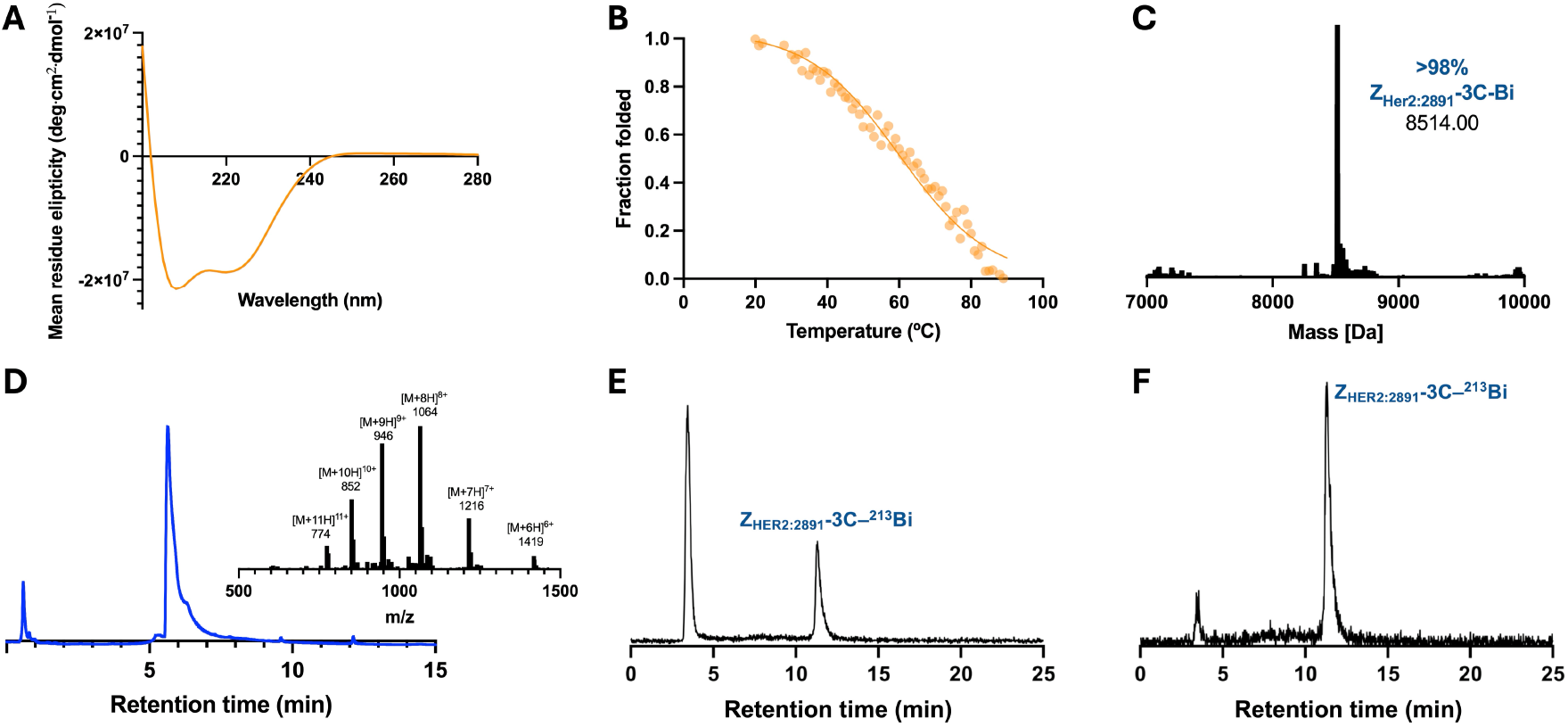
Stability and radiolabeling of Z_HER2:2891_-3C. (A–D) Lyophilized, stored and reconstituted Z_HER2:2891_‐3C samples were reduced and saturated with Bi(III) under optimized conditions, then analyzed by CD spectroscopy (T_m_ = 71 °C) in phosphate buffer, pH 7.5 (A,B), native MS in ammonium acetate buffer, pH 7.0 (C), and LC-MS (chromatogram with m/z inset) using acetonitrile/water with 0.1% formic acid (D), confirming retention of fold and metal‐binding capability. (E,F) Radio‐HPLC analysis (acetonitrile/water with 0.1% TFA) of reconstituted and reduced Z_HER2:2891_‐3C labeled with 1 MBq of ^225^AcCl_3_ generator eluate, shown using the full γ‐window (E) and the ^213^Bi‐specific γ‐window at 440.5 keV (F). The latter indicates a radiochemical purity of >90%. Instrumentation and chromatographic conditions differ between panels (D) and (E,F), resulting in different retention times.

Having confirmed that a lyophilized affibody can be readily reconstituted and analyzed by HPLC, we shipped a lyophilized sample of Z_HER2:2891_-3C from Australia to Germany at ambient temperature to test uptake of the short-lived α-emitter ^213^Bi. After standard reconstitution and reduction with TCEP for 1 h, a ^225^AcCl_3_ stock solution containing the daughter radionuclide ^213^Bi was added and the mixture was analyzed directly by radio-HPLC (Figure 5E,F). ^225^Ac (t_1/2_ = 10 days) decays through a series of short-lived daughter isotopes, mainly ^221^Fr (t_1/2_ = 4.9 min) and ^217^At (t_1/2_ = 32 ms), ultimately forming ^213^Bi (t_1/2_ = 46 min), which itself further mainly decays to ^213^Po and ^209^Pb, before forming stable ^209^Bi. As expected from our initial controls with ^209^Bi, the Z_HER2:2891_-3C–^213^Bi complex produced a distinct and well-resolved peak in the radio-HPLC chromatogram at 11.5 min retention time (Figure 5E,F). When the full γ-window was used, the chromatogram displayed a large injection peak at 4 min, reflecting predominantly unbound radionuclides (e.g., ^221^Fr) and potentially ^213^Bi (Figure 5E). In contrast, restricting the γ‐window to the 440.5 keV line specific for ^213^Bi revealed only a very small injection peak and a dominant peak at 11.5 min (Figure 5F), confirming near‐quantitative complexation of ^213^Bi by the affibody even under harsh, acidic HPLC conditions. To validate these findings, radio-TLC was performed using citrate buffer as the mobile phase for up to five days (Figure S40). Initially, the start position showed a clear signal corresponding to affibody-bound ^213^Bi, while the front contained free radionuclides such as ^221^Fr. After five days, all ^213^Bi at the start had decayed and signals corresponding to longer-lived nuclides remained at the front, consistent with the expected decay behavior.

## Conclusion

In summary, our study demonstrates that a simple triple‐cysteine motif engineered into a helix bundle enables robust, chelator‐free binding of key metals relevant to imaging and therapy, including Bi(III), Pb(II), In(III) and Ga(III). All resulting metal-affibody complexes retain the native fold, exhibit rapid uptake under mild conditions and show high stability against glutathione. Notably, the Bi(III) conjugate displays exceptional resistance to even strong chelators such as EDTA and DTPA, consistent with a buried coordination environment. The ability to lyophilize, ship and reconstitute the engineered affibody without loss of structure or metal‐binding capability supports its practical use in clinical settings. Efficient radiolabeling with the α‐emitter ^213^Bi confirms the suitability of this platform for TAT. Together, these results establish triple‐cysteine affibodies as a promising and operationally simple approach for chelator‐free radiopharmaceutical development.

## Acknowledgements

We are grateful to the Australian Research Council for a Discovery Project (DP230100079) and a Future Fellowship (FT220100010).

