## Supporting Information for "Chelator-Free Radiometal Labeling Inside Engineered Affibodies"

**Materials**

All plasmids encoding an N-terminal $His_6$-tag were obtained from Twist Bioscience, USA. Gastrodenol (bismuth tripotassium dicitrate), glutathione (~98% reduced), tris(hydroxymethyl)aminomethane (Tris) and IPTG were purchased from AK Scientific, USA. Indium(III) chloride hexahydrate, gallium(III) nitrate hydrate, lead(II) nitrate, ammonium acetate (LC-MS grade) and imidazole were purchased from Sigma Aldrich, Australia. TCEP hydrochloride was purchased from AmBeed, USA. Kanamycin was purchased from AG Scientific, USA. Other chemicals and buffer compositions, such as EDTA and DTPA were purchased from Sigma Aldrich, Australia. LC-MS grade solvents were purchased from Fisher Scientific, Australia. Bacterial media components were purchased from Gibco, Thermo Fisher Scientific, USA. HisTrap™ (5 mL) columns were purchased from Cytiva, USA. Pre-cast SDS-PAGE gels (Novex Bis-Tris Plus Mini Protein Gels) were purchased from Invitrogen, Thermo Fisher Scientific, USA. Unstained protein standard, broad range (10-200 KDa) was purchased from New England Biolabs, USA. Amicon®-Ultra centrifugal filters were purchased from Merck Millipore, USA.

**Protein mass spectrometry (MS)**

Intact protein MS

Intact protein analysis was performed on Orbitrap Elite and Orbitrap Fusion™ Tribrid™ mass spectrometers (Thermo Fisher Scientific, USA) connected to a Thermo Fisher Scientific UltiMate 3000 HPLC system equipped with a ZORBAX 300SB-C3, 3.5 µm, 4.6 × 50 mm HPLC column (Agilent Technologies, USA). 10–20 µM protein samples were injected using a 500 µL/min linear gradient of solvent A (0.1% (*v*/*v*) formic acid in water) and solvent B (0.1% (*v*/*v*) formic acid in acetonitrile), ramping solvent B from 5% at the start to 80% after 12 min. The ion source was H-ESI with a static spray voltage set to 3500 V in positive ion mode. The sheath, auxiliary and sweep gasses were set to 50 Arb, 10 Arb and 1 Arb, respectively. The ion transfer tube temperature was set to 325 °C and the vaporizer temperature was set to 350 °C. The resolution of the Orbitrap MS detector was set to 120000, and the scan range was set to 500–4000 m/z. Protein mass was determined by deconvolution using the program Xcalibur 3.0.63 (Thermo Fisher Scientific, USA).

Native MS with prior buffer exchange

Samples for native MS were exchanged into 100 mM $NH_4OAc$ (pH 7) in a 3 kDa MWCO centrifugal filter unit (Amicon®-Ultra 0.5) prior to analysis. Native protein analysis was performed on an Orbitrap Fusion™ Tribrid™ mass spectrometer (Thermo Fisher Scientific,

USA) connected to a Thermo Fisher Scientific UltiMate 3000 HPLC system. 10–20 μM protein samples were injected using a 5 minute 100 μL/min isocratic elution mode in 100 mM $NH_4OAc$ (pH 7). The ion source was H-ESI with a static spray voltage set to 3000 V in positive ion mode. The sheath, auxiliary and sweep gasses were set to 25 Arb, 5 Arb and 0 Arb, respectively. The ion transfer tube temperature was set to 275 °C and the vaporizer temperature was set to 50 °C. The MS detector was an Orbitrap with the resolution set to 120000, and the scan range was set to 500–4000 m/z. Native protein mass was determined by deconvolution using the program Xcalibur 3.0.63 (Thermo Fisher Scientific, USA).

Alternative native MS with prior buffer exchange

An alternative native MS method was performed on a Waters Synapt G2-Si HDMS qTOF mass spectrometer connected to a Waters Acquity UPLC I Class plus LC unit. 10–20 μM protein samples were injected using a 3 minute 100 μL/min isocratic elution mode in 100 mM $NH_4OAc$ (pH 7). The mass spectrometer was operated in positive, full MS, resolution and TOF modes. Capillary voltage and cone voltage were set to 1.5 kV and 30 V, respectively. The source temperature was set to 120 °C and the desolvation temperature was set to 350 °C. Leucine enkephalin was used as the Lockspray reference compound.

**Affibody expression and purification**

$Z_{HER2:2891}$ and $Z_{HER2:2891}$-3C

Affibody sequences with N-terminal $His_6$-tag (Table S1) were cloned into a pET-29b(+) expression vector (Twist Bioscience, USA) and transformed into *E. coli* BL21 (DE3) cells.[1] Isolated single colonies were grown in Luria Broth (LB) overnight, transferred to 1 L LB and grown at 37 °C with gentle shaking until the optical density (OD) reached 0.6–0.8. This was followed by induction of overexpression using 1 mM IPTG for 18–20 h at 25 °C. The *E. coli* cells were harvested (centrifuged at 5000 × g, 30 minutes, 4 °C) and resuspended in lysis buffer (20 mM Tris-HCl, pH 8) for 30 minutes on ice with gentle shaking. Cells were lysed by sonication (Omni Sonic Ruptor 400 Ultrasonic homogenizer) three times at 50% power for 30 seconds on ice, before centrifugation at 17,000 × g for 30 minutes to separate cell debris. The soluble fraction was then loaded onto a 5 mL HisTrap™ HP column (Cytiva) which had been equilibrated with binding buffer (20 mM Tris-HCl, pH 8.0, 300 mM NaCl, 15 mM imidazole). The column was then washed with 5 column volumes of the binding buffer. Protein was eluted using 5 column volumes of elution buffer (20 mM Tris-HCl, pH 8.0, 200 mM NaCl, 300 mM imidazole) and the protein sample was exchanged into and concentrated in storage buffer (10 mM Tris-HCl, 150 mM NaCl, pH 7.5) using a 3 kDa MWCO centrifugal filter unit (Amicon®-

Ultra 15). The protein solution was aliquoted, and flash frozen in liquid nitrogen for long term storage at –80 °C.

$Z_{TNF\text{-}alpha1}$ and $Z_{TNF\text{-}alpha1}$-3C

Affibody sequences with N-terminal $His_6$-tag (Table S1) were cloned into a pET-29b(+) expression vector (Twist Bioscience, USA) and transformed into *E. coli* BL21 (DE3) cells. The expression protocol was followed as described by Kronqvist et al. with minor changes.[2] Isolated single colonies were grown in Luria Broth (LB) overnight, transferred to 1 L LB and grown at 37 °C with gentle shaking until the OD reached 0.8–1.0. This was followed by induction of overexpression using 1 mM IPTG for 4 h at 37 °C. The *E. coli* cells were harvested (centrifuged at 5000 × g, 30 minutes, 4 °C) and resuspended in binding buffer (PBS, pH 7.4, 30 mM imidazole) for 30 minutes on ice with gentle shaking. Cells were lysed by sonication (Omni Sonic Ruptor 400 Ultrasonic homogenizer) three times at 50% power for 30 seconds on ice, before centrifugation at 17,000 × g for 30 minutes to separate cell debris. The soluble fraction was then loaded onto a 5 mL HisTrap™ HP column (Cytiva) which had been equilibrated with binding buffer. The column was then washed with 5 column volumes of the binding buffer. Protein was eluted using 5 column volumes of elution buffer (PBS, pH 7.4, 200 mM imidazole), followed by exchange into and concentration in storage buffer (10 mM Tris-HCl, 150 mM NaCl, pH 7.5) using a 3 kDa MWCO centrifugal filter unit (Amicon®-Ultra 15). The protein solution was aliquoted, and flash frozen in liquid nitrogen for long term storage at –80 °C.

Uniformly $^{15}N$-labelled $Z_{HER2:2891}$-3C

Cells were grown at 37 °C until the OD reached 0.6–0.8 in LB. Cells were centrifuged (5000 × g, 30 minutes, 4 °C) and resuspended in minimal medium (50 mM $NaHPO_4$, 25 mM $KH_2PO_4$, 10 mM NaCl, pH 8.0) supplemented with 5 mM $MgSO_4$, 0.2 mM $CaCl_2$, 0.25% metal mix, $^{15}NH_4Cl$ (1g/L) as the only nitrogen source, and 1% glucose. Cells were grown in minimal medium at 37 °C for 30 minutes before induction using 1 mM IPTG and overexpression for 18–20 h at 25 °C. Cell lysis, extraction and purification were performed as described above. The $^{15}N$-labelled affibody was dialysed overnight into NMR buffer (20 mM MES, pH 6.5, 150 mM NaCl), flash frozen in liquid nitrogen and stored at –80 °C.

All proteins were characterized and their correct mass confirmed by SDS-PAGE and intact protein MS (positive ion mode).

**Metal uptake reaction**

Optimisation

Initial attempts to reduce and modify the affibodies were adapted from our previously established methods whereby we first reduce the protein at a temperature 8 – 10 °C below their denaturation midpoint ($T_m$) under acidic conditions in the presence of an excess of the reducing agent TCEP for up to 60 minutes prior to addition of 5 equivalents of the relevant metal.[3] With these methods proving successful, we then sought out milder reduction conditions at 25 °C and/or with a pH of 7.5.

Optimised metal uptake reaction

Affibodies (100 µM in 10 mM Tris-HCl, 150 mM NaCl, pH 7.5) were reduced at 25 °C in the presence of 50 mM TCEP (pH 7.5) for a maximum duration of 60 minutes. Five equivalents of the metal salts gastrodenol (for bismuth), $InCl_3$ hexahydrate, $Ga(NO_3)_3$ hydrate or $Pb(NO_3)_2$ were added to the reaction solution from their respective stocks (50 mM in water), and immediately vortexed for 60 seconds. Following the reaction, the solution was quickly centrifuged, and the supernatant was collected for further analysis. For quantifying metal uptake by native MS, the supernatant was exchanged into 100 mM $NH_4OAc$ (pH 7).

**Circular dichroism (CD) spectroscopy**

Secondary structures of the affibodies were assessed by circular dichroism using a Chirascan spectropolarimeter from Applied Photophysics equipped with a temperature control module using a 0.1 cm path-length cuvette. Each CD spectrum was averaged over two scans and the baseline correction was done by blank subtraction. Individual affibodies were diluted in 20 mM phosphate buffer, pH 7.5. Scans were carried out at 25 °C covering a range of 200–280 nm with a 1 nm step size and 1 nm bandwidth. Thermal denaturation experiments were monitored at 222 nm over a temperature range of 20–90 °C (1 °C/min heating rate). Temperature-dependent CD data were fitted to a two-state unfolding model (Boltzmann sigmoidal equation) and plotted in GraphPad Prism 10 (Dotmatics, USA) to obtain the denaturation midpoint ($T_m$).

**Evaluating the stability of metal-bound affibodies**

Affibodies were reduced and saturated with either Bi(III), In(III), Ga(III) or Pb(II) as per the optimized protocol to obtain >98% metal saturation, prior to buffer exchange into 100 mM $NH_4OAc$ (pH 7). Samples were then stored at 4 °C for up to seven days and analysed via native MS at different time points to determine the stability of the metal-bound affibody.

**Glutathione (GSH) competition assay**

At first, reduction followed by uptake reactions with Bi(III), In(III), Ga(III) and Pb(II) was performed with the affibody $Z_{Her2:2891}$-3C as per the optimized protocol with the addition of 1.2 equivalents of metal to obtain >98% metal saturation. The metal-bound affibody samples were exchanged into 100 mM $NH_4OAc$ (pH 7). A reduced glutathione (GSH) stock was prepared in 100 mM $NH_4OAc$ and adjusted to pH 7. Competition experiments were performed with 1, 10, 20, 50 and 100 equiv. of GSH with respect to the concentrations of the metal-bound affibodies (50 μM) for 1 h at 25 °C. Native MS of the affibodies was performed in 100 mM $NH_4OAc$. Conservation of metal-bound affibodies (%) was plotted as a function of increasing equivalents of GSH in GraphPad Prism 10 (Dotmatics, USA). One-phase exponential decay was used as the fitting function.

The $Z_{Her2:2891}$-3C-Bi in the presence of 100 equiv. GSH sample was stored at 4 °C for up to 14 days and the stability of the metal-bound affibody in the presence of GSH was monitored by native MS over time. Metal-bound affibodies (%) were plotted as a function of time (days) in GraphPad Prism 10 (Dotmatics, USA). One-phase exponential decay was used as the fitting function.

**Ethylenediaminetetraacetic acid (EDTA) competition assay**

At first, reduction followed by uptake reactions with Bi(III), In(III), Ga(III) and Pb(II) was performed with the affibody $Z_{Her2:2891}$-3C or only Bi(III) with the affibody $Z_{TNF\text{-}alpha1}$-3C as per the optimized protocol with the addition of 1.2 equivalents of metal to obtain >98% metal saturation. The metal-bound affibody samples were exchanged into 100 mM $NH_4OAc$ (pH 7). An EDTA stock was prepared in 100 mM $NH_4OAc$ and adjusted to pH 7. Competition experiments were performed with 1, 10, 20, 50, and 100 equiv. of EDTA with respect to the concentrations of the metal-bound affibodies (50 μM) for 1 h at 25 °C. Native MS of the affibodies was performed in 100 mM $NH_4OAc$. Conservation of metal-bound affibodies (%) was plotted as a function of increasing equivalents of EDTA in GraphPad Prism 10 (Dotmatics, USA). One-phase exponential decay was used as the fitting function.

The $Z_{Her2:2891}$-3C-Bi in the presence of 100 equiv. EDTA sample was stored at 4 °C for up to 14 days and the stability of the metal bound affibody in the presence of EDTA was monitored by native MS overtime. Conservation of metal-bound affibodies (%) was plotted as a function of time (days) in GraphPad Prism 10 (Dotmatics, USA). One-phase exponential decay was used as the fitting function.

Another example of the $Z_{Her2:2891}$-3C-Bi in the presence of 100 equiv. EDTA sample was also analysed via the alternative native MS method described above.

Another example of $Z_{Her2:2891}$-3C-Bi was incubated for 10 minutes at 100 °C in the presence of 0 equiv. or 100 equiv. EDTA in 100 mM $NH_4OAc$ pH 7. Native MS of the affibodies was performed in 100 mM $NH_4OAc$. Metal bound affibodies (%) were represented as a bar graph in GraphPad Prism 10 (Dotmatics, USA).

**Diethylenetriaminepentaacetic acid (DTPA) competition assay**

At first, reduction followed by uptake reactions with Bi(III) was performed with the affibody $Z_{Her2:2891}$-3C as per the optimized protocol with the addition of 1.2 equivalents of metal to obtain >98% metal saturation. The metal-bound affibody samples were exchanged into 100 mM $NH_4OAc$ (pH 7) buffer. A DTPA stock was prepared in 100 mM $NH_4OAc$ and adjusted to pH 7. Competition experiments were performed with 1, 10, 20, 50 and 100 equiv. of DTPA with respect to the concentrations of the metal-bound affibodies (50 μM) for 1 h at 25 °C. Native MS of the affibodies was performed in 100 mM $NH_4OAc$. Conservation of metal bound-affibodies (%) was plotted as a function of increasing equivalents of DTPA in GraphPad Prism 10 (Dotmatics, USA). One-phase exponential decay was used as the fitting function.

**Computational modelling**

The amino acid sequence of $Z_{Her2:2891}$ was used as input to generate a structural model using ColabFold v1.5.4:AlphaFold2.[4] This structure was imported into BIOVIA Discovery Studio Visualiser 2025, where amino acids A12, L34 and S41 were exchanged to cysteine residues. One bismuth(III) atom was introduced and connected to the three reduced cysteines. The charges on the cysteine residues and bismuth atom were neutralised and the distance constraints based on known tris(L-cysteinato-*S*)-bismuth(III) monohydrate (CCDC entry: CIYPIK) coordination geometry were applied.[5] Geometry optimisation was then performed using the CHARMM force field. Figures were visualised using ChimeraX software.[6]

**NMR spectroscopy**

All NMR spectra were recorded at 25 °C using an 800 MHz Bruker Avance NMR spectrometer equipped with a cryoprobe. [$^{15}$N,$^{1}$H]-HSQC spectra were recorded in a 3 mm NMR tube. The affibody sample (400 μM in 20 mM MES, pH 6.5, 150 mM NaCl) was exposed to 20 mM TCEP (pH 7.5) for 60 minutes at 25 °C and an NMR spectrum was collected of the reduced uniformly $^{15}N$-labeled $Z_{Her2:2891}$-3C in NMR buffer (20 mM MES, pH 6.5, 150 mM NaCl, 10% $D_2O$). For Bi(III) uptake, gastrodenol (bismuth source) was titrated in and an [$^{15}N$, $^{1}H$]-HSQC spectrum was collected at each stage with the same parameters as the reduced apo-affibody. The spectrum representing the equivalent protein-metal complex was used.

**Experiments with lyophilised samples**

Lyophilisation

100 μL of 100 μM affibody sample in 10 mM Tris-HCl, 150 mM NaCl, pH 7.5 was flash frozen in liquid nitrogen and lyophilised. Lyophilised samples were stored at room temperature until needed.

CD spectroscopy

The lyophilised sample was redissolved in 100 μL MilliQ water. 50 uL of 100 mM TCEP, pH 7, was added and the sample was incubated at 25 ºC for 60 minutes. 5 equiv. gastrodenol was added and the sample was immediately vortexed. CD spectroscopy was performed as described above.

Native MS

The lyophilised sample was redissolved in 100 μL MilliQ water. 50 uL of 100 mM TCEP, pH 7, was added and the sample was incubated at 25 ºC for 60 minutes. 5 equiv. gastrodenol was added and the sample was immediately vortexed. Native MS was performed as described above (alternative native MS method with prior buffer exchange).

**Analytical LCMS**

Analytical LC-MS was conducted on an Agilent HPLC-MS (1260/6120) equipped with a reverse phase column (Poroshell 120 EC-C18, 2.7 μm, 3.0 × 50 mm) held at 30 ºC, where elution was monitored via UV absorbance at 280 nm and m/z ratios were confirmed via mass spectrometry. A binary gradient system of 5-95% ACN:$H_2O$ with 0.1% formic acid (*v/v*) over 15 minutes with a flow rate of 0.3 mL/min was used. The gradient started at 5% ACN, followed by a gradual increase to 90% over 10 minutes where it was held for a further five minutes.

**Radiolabelling experiments with $^{213}$Bi**

Synthesis of $Z_{Her2:2891}$-3C-$^{213}$Bi

$^{225}$Ac was obtained from the Joint Research Centre Karlsruhe. 100µL of a 100 mM solution of $Z_{Her2:2891}$-3C (reconstituted from a lyophilised sample) was mixed with 50 µL of a 50 mM TCEP solution, pH 8.1. After 60-minute incubation at 25 °C, the mixture was purified via size-exclusion chromatography filter units (Amicon Ultra Centrifugal Units with 3 kDa MWCO cut-off). The purified solution was mixed with 1 µL (1 MBq) $^{225}AcCl_3$-solution (provided by JRC Karlsruhe) at 25 °C and HPLC as well as TLC were performed immediately after the addition of radioactivity.

HPLC with $Z_{Her2:2891}$-3C-$^{213}$Bi

The HPLC measurement was performed using a Jasco HPLC system (UV-975 UV detector and PU-980 HPLC Pump). A Phenomenex Aeris WIDEPORE 3.6u XB-C18 250 x 4.60 mm column was used (00G-4482-E0). The following gradient was used: Solvent A acetonitrile + 0.1% TFA, solvent B water + 0.1% TFA with a flow set at 0.8 mL/min: 0 min to 25 min 5%-95% solvent A. The temperature was not controlled. 20 µL with 200-300 kBq were injected using a manual injector per measurement.

An ElysiaRaytest Gabi Nova $LaBr_3$ (Straubenhardt, Germany) detector was used with either a 0-2000 keV window or 420-500 keV window. The signals were analyzed using the software Gina X.

Thin layer chromatography with $Z_{Her2:2891}$-3C-$^{213}$Bi

TLC was performed using the TLC scanner miniGita from Raytest. The mobile phase was citrate-dextrose solution (ACD) C3821-50ML (pH 4.5-5.5) from Sigma-Aldrich. The stationary phase was C18 silica gel. A 1 µL spot of synthesized $Z_{Her2:2891}$-3C-$^{213}$Bi was placed on the TLC plate and after 2/3 of the plate was developed, the plate was analyzed with the TLC scanner at t = 0 min and t = 5 days.

**Table S1**. Sequences of affibody constructs used in this study (mutations highlighted in orange, $His_6$-tag in blue).

| Affibody | Sequence |
|---|---|
| $Z_{HER2:2891}$ | MMGSSHHHHHHLQAEAKYAKEMRNAYWEIALLPNLTNQQKRAFIRKLY DDPSQSSELLSEAKKLNDSQAPK |
| $Z_{HER2:2891}$-3C | MMGSSHHHHHHLQAEAKYAKEMRNCYWEIALLPNLTNQQKRAFIRKCY DDPSQCSELLSEAKKLNDSQAPK |
| $Z_{TNF\text{-}alpha1}$ | MGSSHHHHHHLQVDNKFNKENIAAMTEITRLPNLNPYQRAAFIWSLSDD PSQSANLLAEAKKLNDAQAPK |
| $Z_{TNF\text{-}alpha1}$-3C | MMGSSHHHHHHLQVDNKFNKENIACMTEITRLPNLNPYQRAAFIWSCSD DPSQCANLLAEAKKLNDAQAPK |

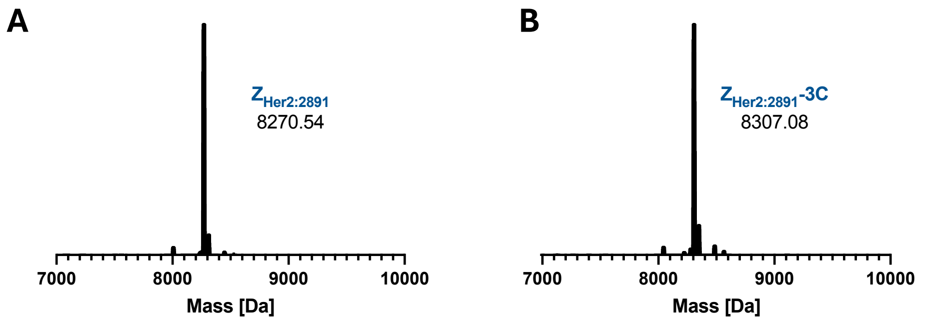


**Figure S1:** Intact MS of $Z_{HER2:2891}$ wildtype (A) and $Z_{HER2:2891}$-3C (B) after recombinant expression in *E. coli.*

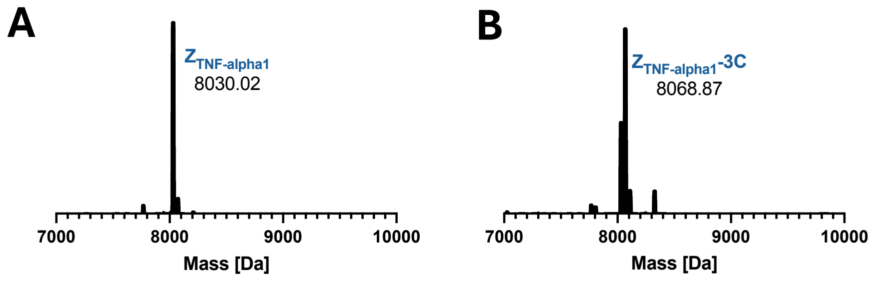


**Figure S2:** Intact MS of $Z_{TNF\text{-}alpha1}$ wildtype (A) and $Z_{TNF\text{-}alpha1}$-3C (B) after recombinant expression in *E. coli.*

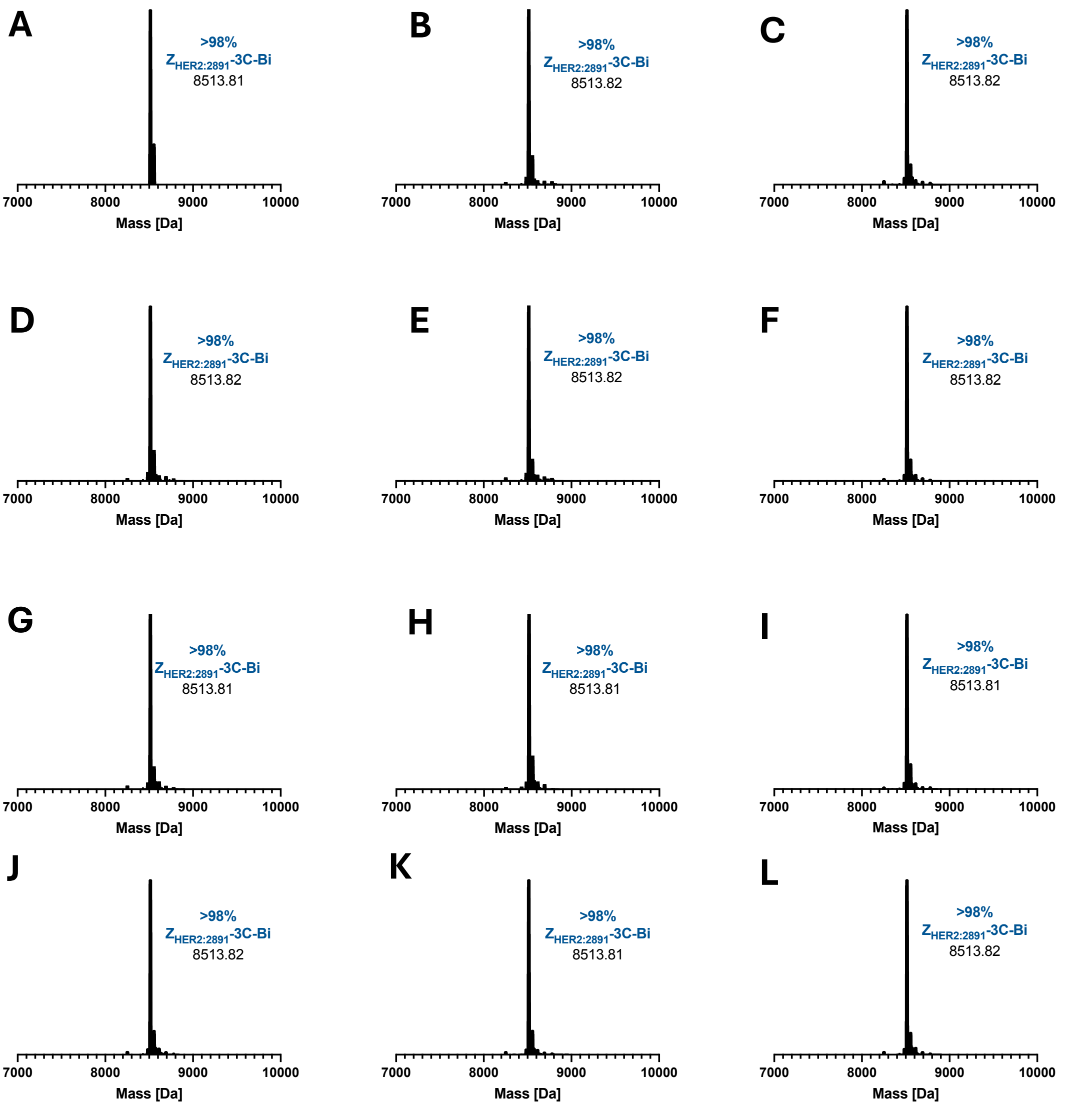


**Figure S3:** Uptake of Bi(III) by $Z_{HER2:2891}$-3C in triplicate determined by native MS (deconvoluted) after incubation at 25 °C in 20 mM Tris, pH 7.5, 150 mM NaCl, 50 mM TCEP for 5 min (A-C), 15 min (D-F), 30 min (G-I) and 60 min (J-L).

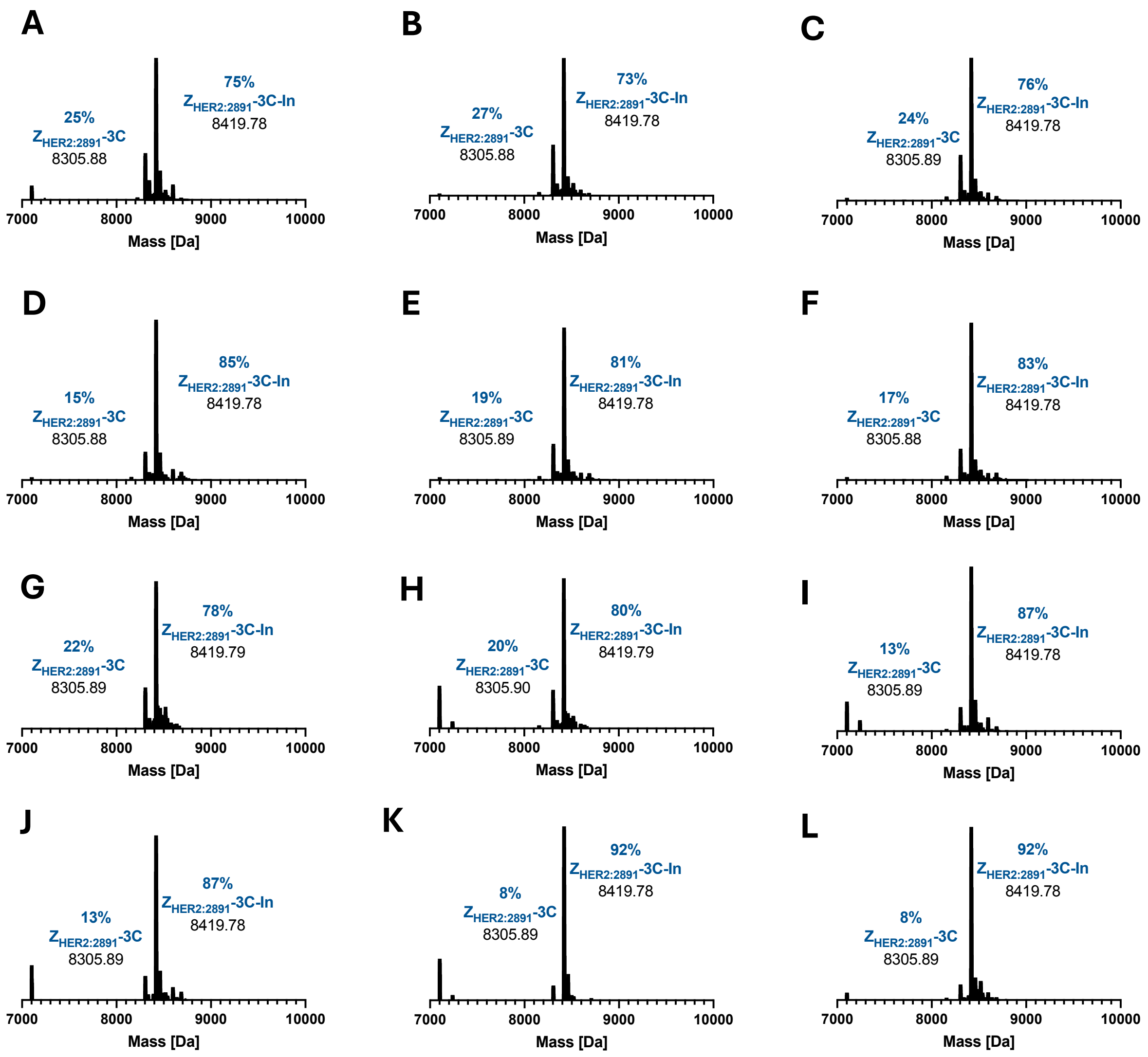


**Figure S4:** Uptake of In(III) by Z$_{HER2:2891}$-3C in triplicate determined by native MS (deconvoluted) after incubation at 25 °C in 20 mM Tris, pH 7.5, 150 mM NaCl, 50 mM TCEP for 5 min (A-C), 15 min (D-F), 30 min (G-I) and 60 min (J-L).

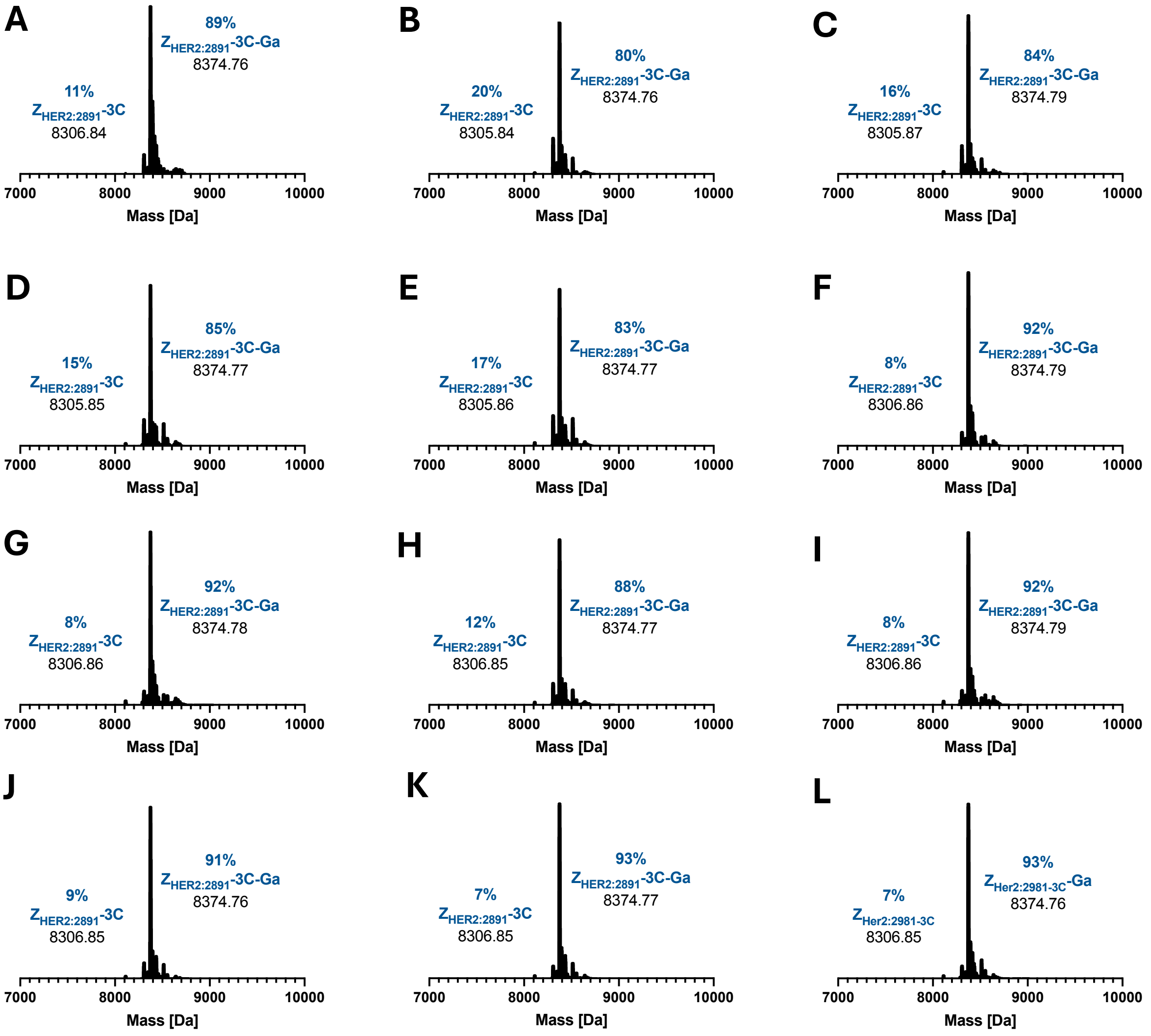


**Figure S5:** Uptake of Ga(III) by $Z_{HER2:2891}$-3C in triplicate determined by native MS (deconvoluted) after incubation at 25 °C in 20 mM Tris, pH 7.5, 150 mM NaCl, 50 mM TCEP for 5 min (A-C), 15 min (D-F), 30 min (G-I) and 60 min (J-L).

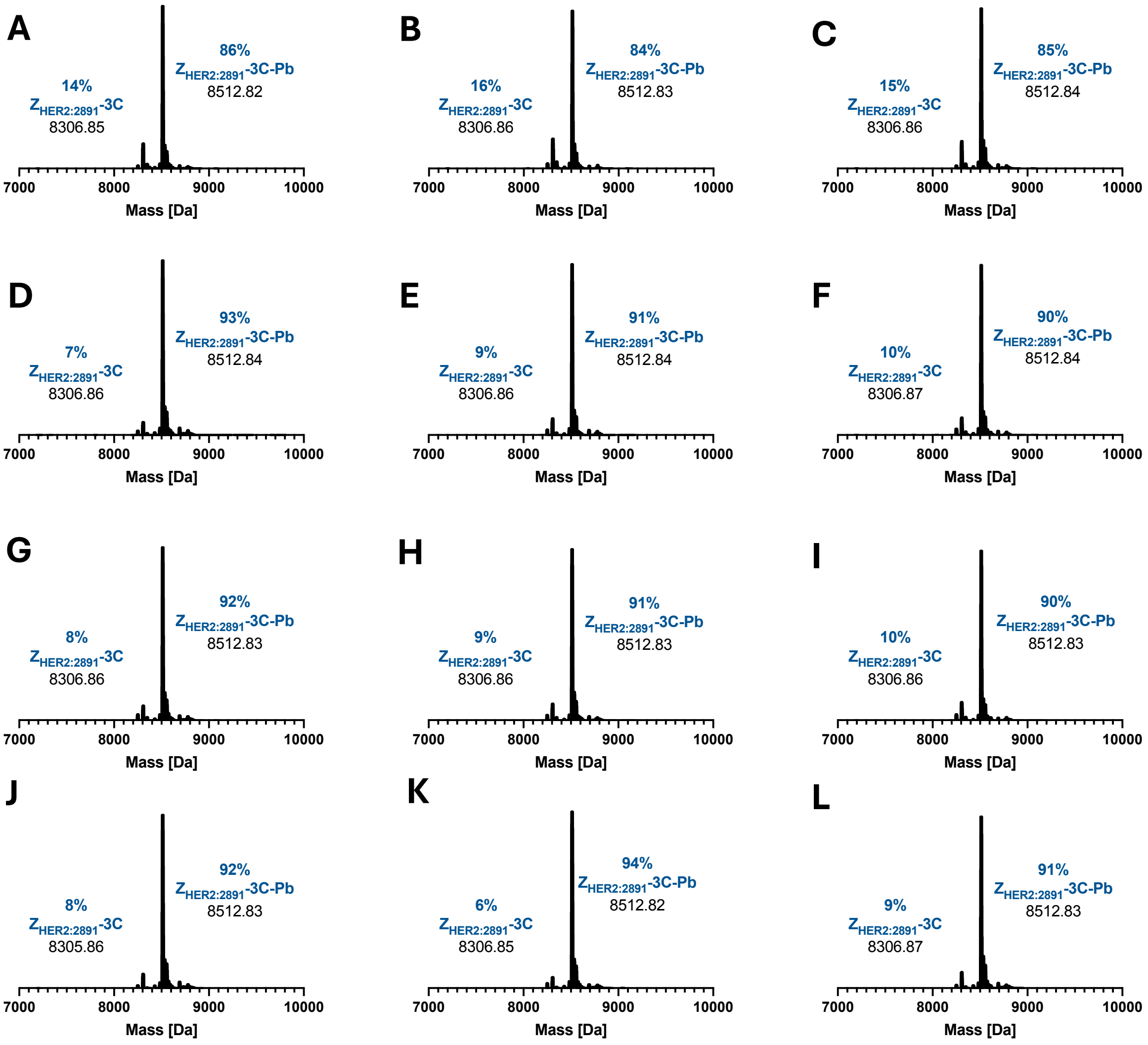


**Figure S6:** Uptake of Pb(II) by $Z_{HER2:2891}$-3C in triplicate determined by native MS (deconvoluted) after incubation at 25 °C in 20 mM Tris, pH 7.5, 150 mM NaCl, 50 mM TCEP for 5 min (A-C), 15 min (D-F), 30 min (G-I) and 60 min (J-L).

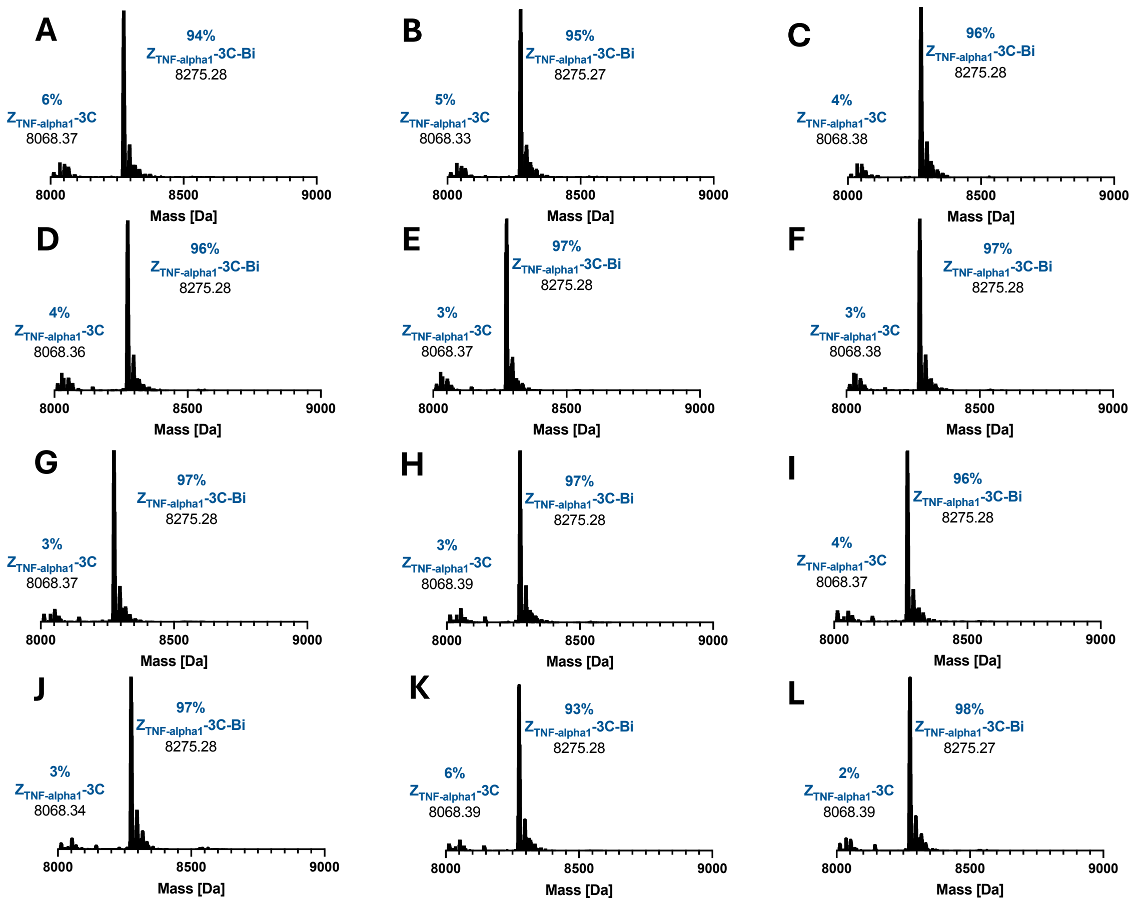


**Figure S7:** Uptake of Bi(III) by $Z_{TNF\text{-}alpha1}$-3C in triplicate determined by native MS (deconvoluted) after incubation at 25 °C in 20 mM Tris, pH 7.5, 150 mM NaCl, 50 mM TCEP for 5 min (A-C), 15 min (D-F), 30 min (G-I) and 60 min (J-L).

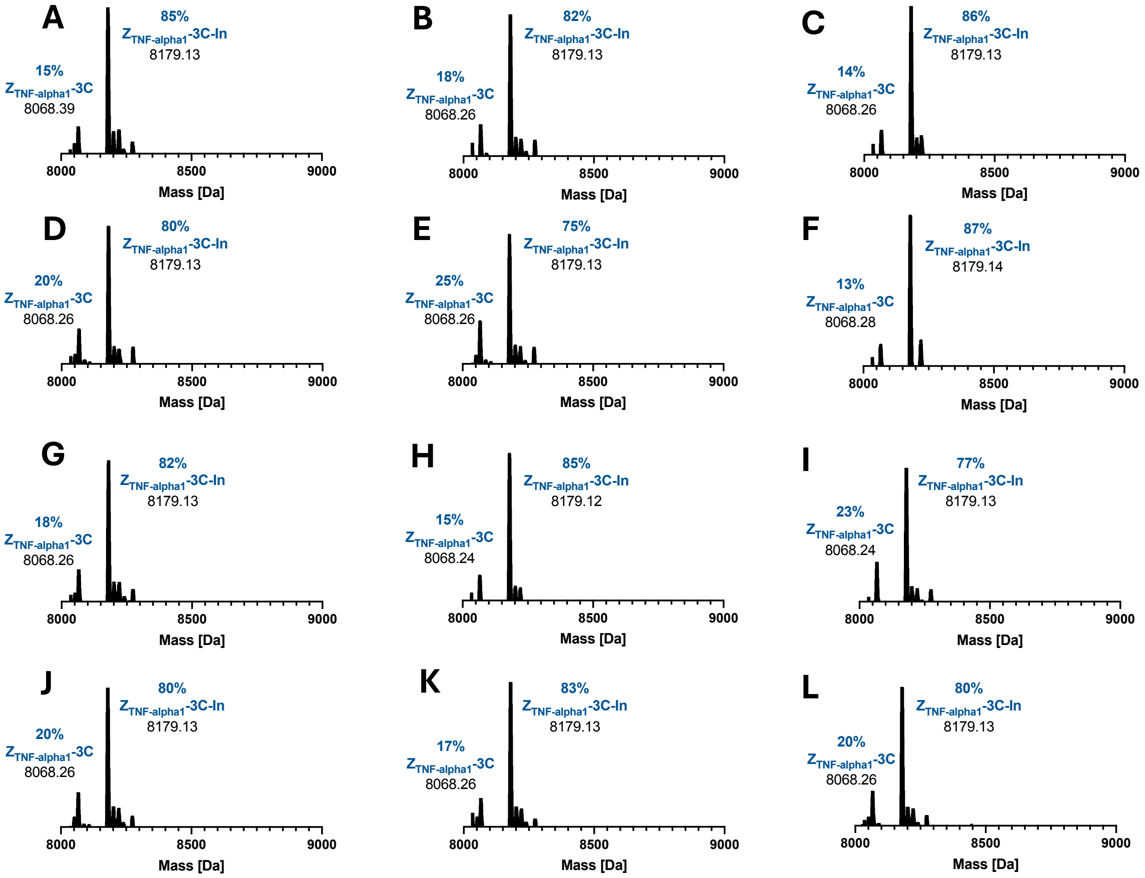


**Figure S8:** Uptake of In(III) by $Z_{TNF\text{-}alpha1}$-3C in triplicate determined by native MS (deconvoluted) after incubation at 25 °C in 20 mM Tris, pH 7.5, 150 mM NaCl, 50 mM TCEP for 5 min (A-C), 15 min (D-F), 30 min (G-I) and 60 min (J-L).

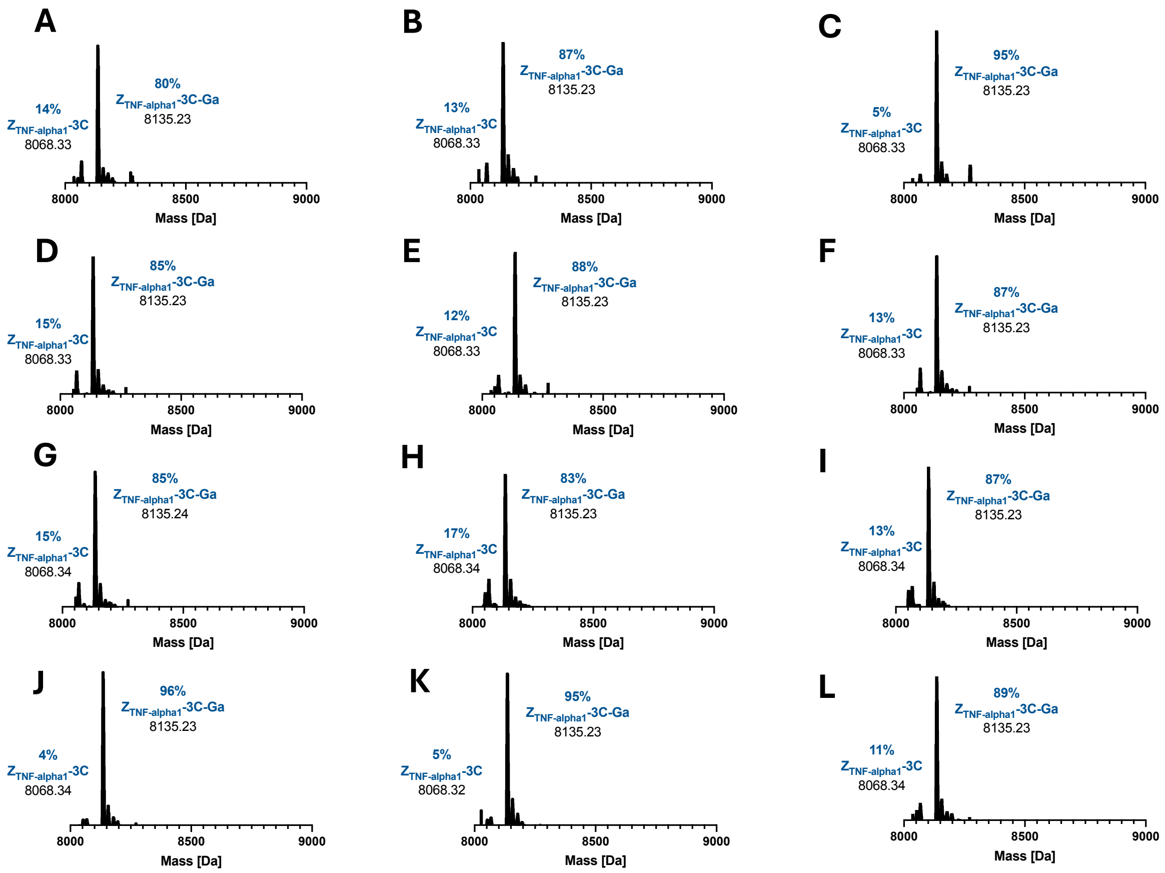


**Figure S9:** Uptake of Ga(III) by $Z_{TNF\text{-}alpha1}$-3C in triplicate determined by native MS (deconvoluted) after incubation at 25 °C in 20 mM Tris, pH 7.5, 150 mM NaCl, 50 mM TCEP for 5 min (A-C), 15 min (D-F), 30 min (G-I) and 60 min (J-L).

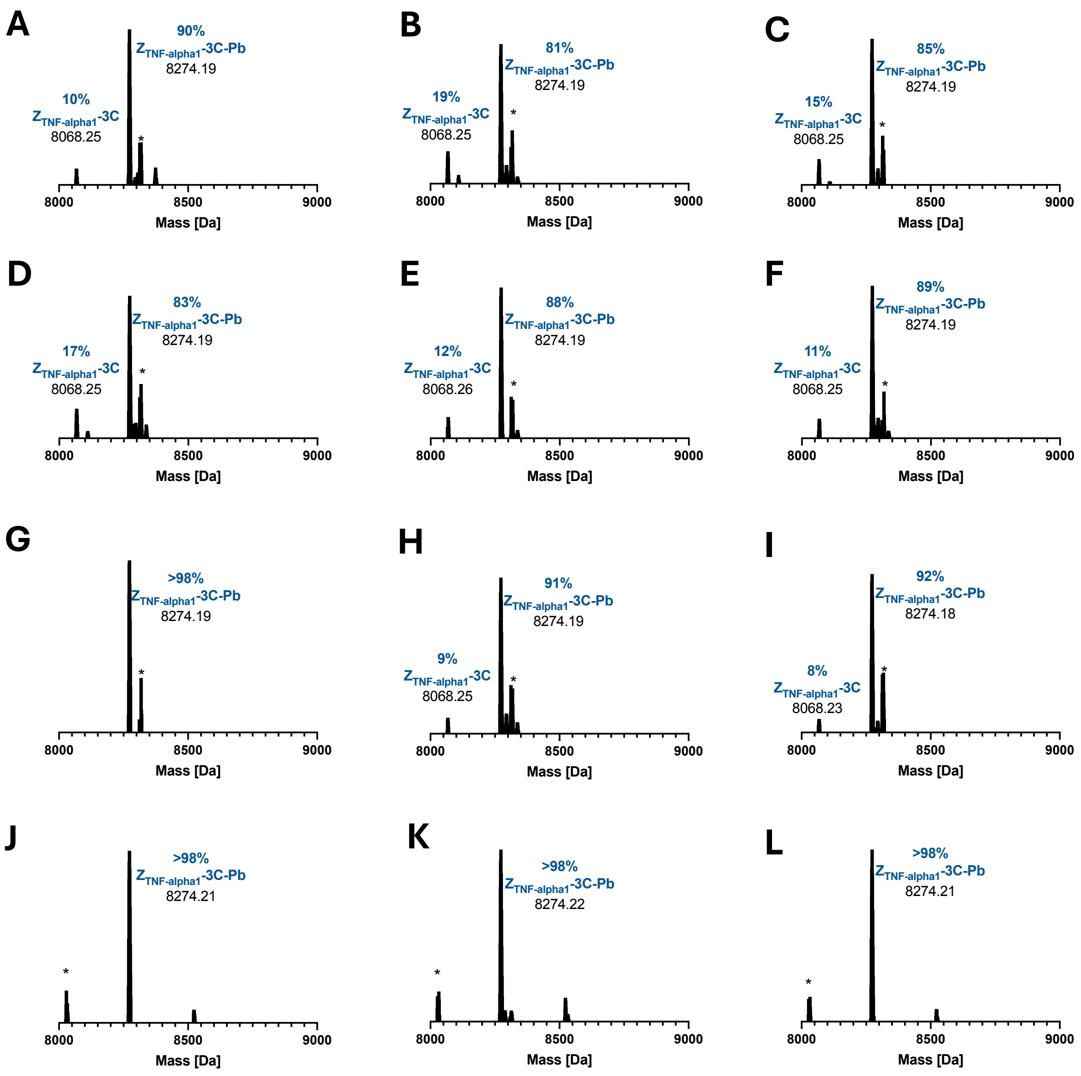


**Figure S10:** Uptake of Pb(II) by $Z_{TNF\text{-}alpha1}$-3C in triplicate determined by native MS (deconvoluted) after incubation at 25 °C in 20 mM Tris, pH 7.5, 150 mM NaCl, 50 mM TCEP for 5 min (A-C), 15 min (D-F), 30 min (G-I) and 60 min (J-L). Minor impurities are indicated by an asterisk (*).

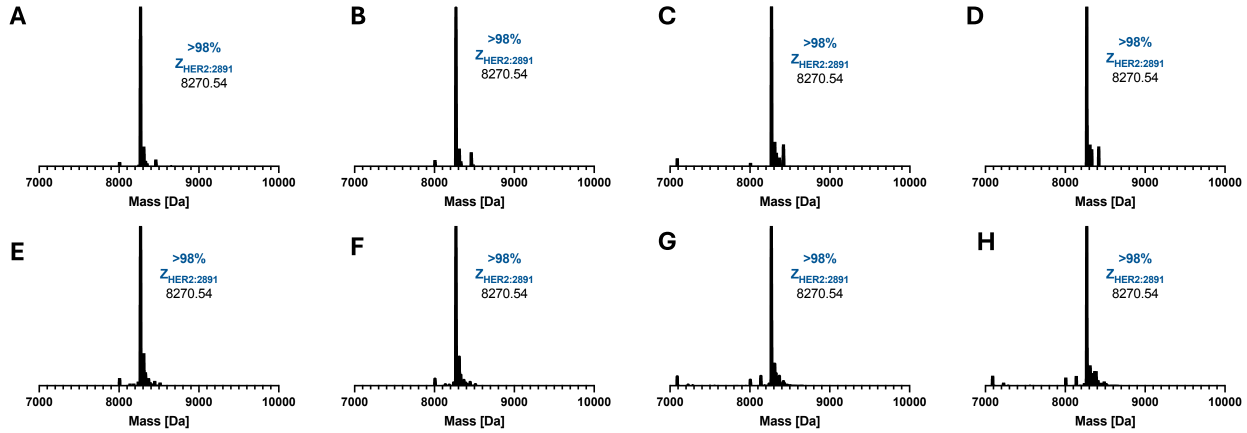


**Figure S11:** Native MS (deconvoluted) of the $Z_{HER2:2891}$ wildtype (with $His_6$-tag) in 100 mM ammonium acetate pH 7.0 indicating no uptake of Bi(III) (A, B), In(III) (C, D), Ga(III) (E, F) or Pb(II) (G, H) after exposure to gastrodenol (5 equiv.), indium(III) chloride (5 equiv.), gallium(III) nitrate hydrate (5 equiv.) or lead(II) nitrate (5 equiv.) under the optimised conditions.

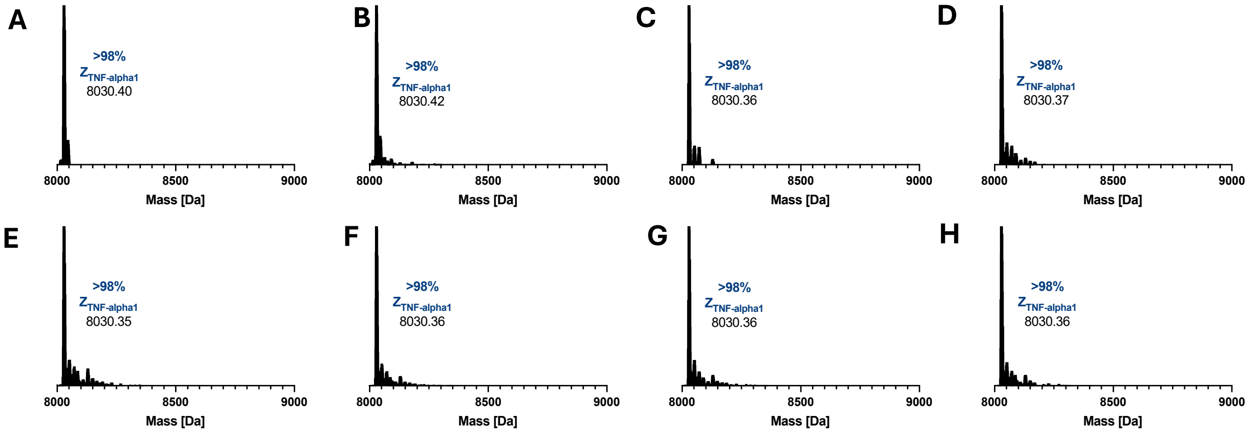


**Figure S12:** Native MS (deconvoluted) of the $Z_{TNF\text{-}alpha1}$ wildtype (with $His_6$-tag) in 100 mM ammonium acetate pH 7.0 indicating no uptake of Bi(III) (A, B), In(III) (C, D), Ga(III) (E, F) or Pb(II) (G, H) after exposure to gastrodenol (5 equiv.), indium(III) chloride (5 equiv.), gallium(III) nitrate hydrate (5 equiv.) or lead(II) nitrate (5 equiv.) under the optimised conditions.

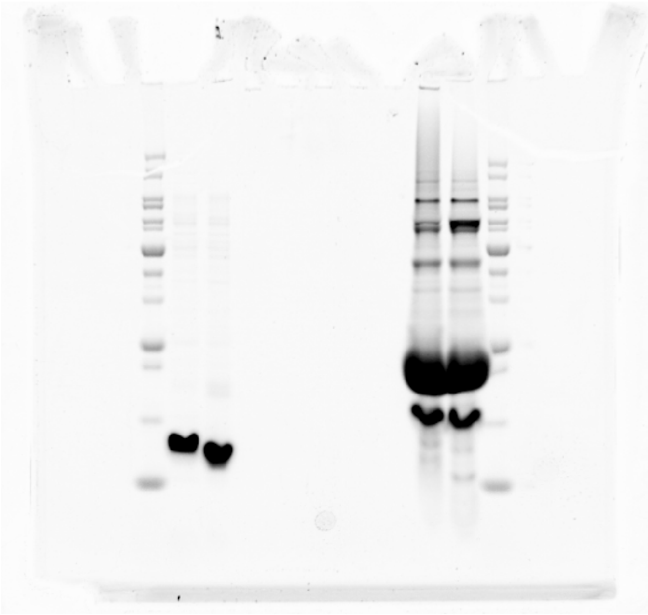

**Figure S13:** Uncropped SDS-PAGE image of data shown (lanes 3-5) in Figure 2G.

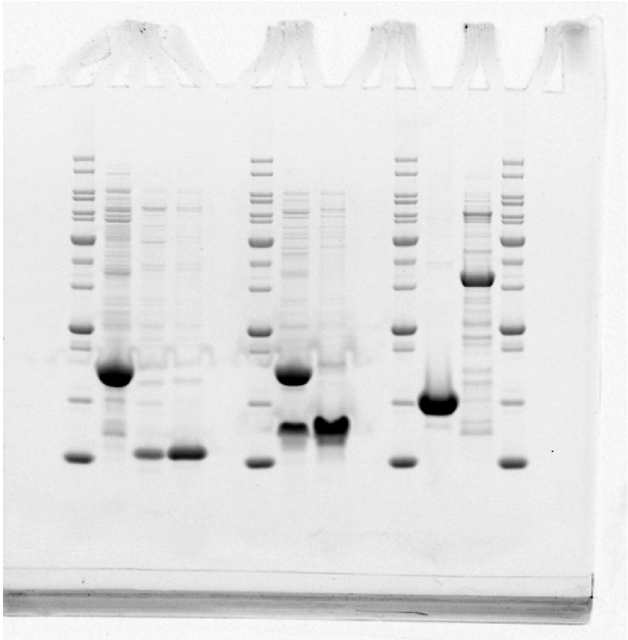

**Figure S14:** Uncropped SDS-PAGE image of data shown (lanes 2,4-5) in Figure 2H.

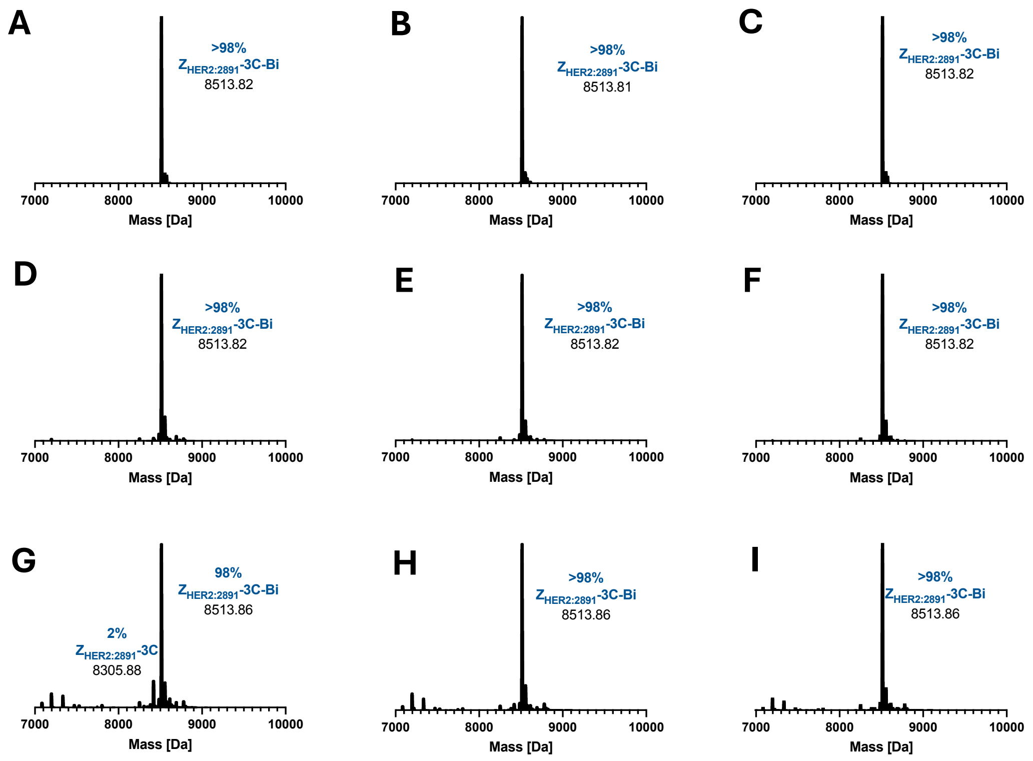


**Figure S15:** Native MS (deconvoluted) of Bi(III)-bound $Z_{HER2:2891}$-3C in triplicate stored in 100 mM ammonium acetate pH 7 at 4 °C for 0 days (A-C), 1 day (D-F) and 7 days (G-I).

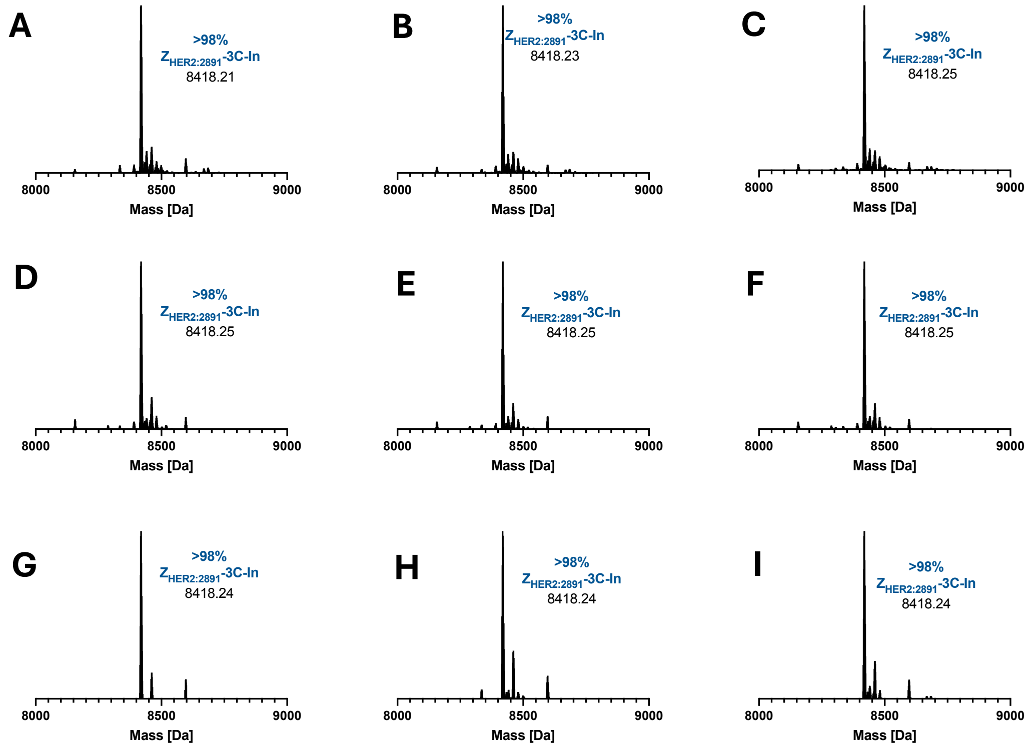


**Figure S16:** Native MS (deconvoluted) of In(III)-bound $Z_{HER2:2891}$-3C in triplicate stored in 100 mM ammonium acetate pH 7 at 4 °C for 0 days (A-C), 1 day (D-F) and 7 days (G-I).

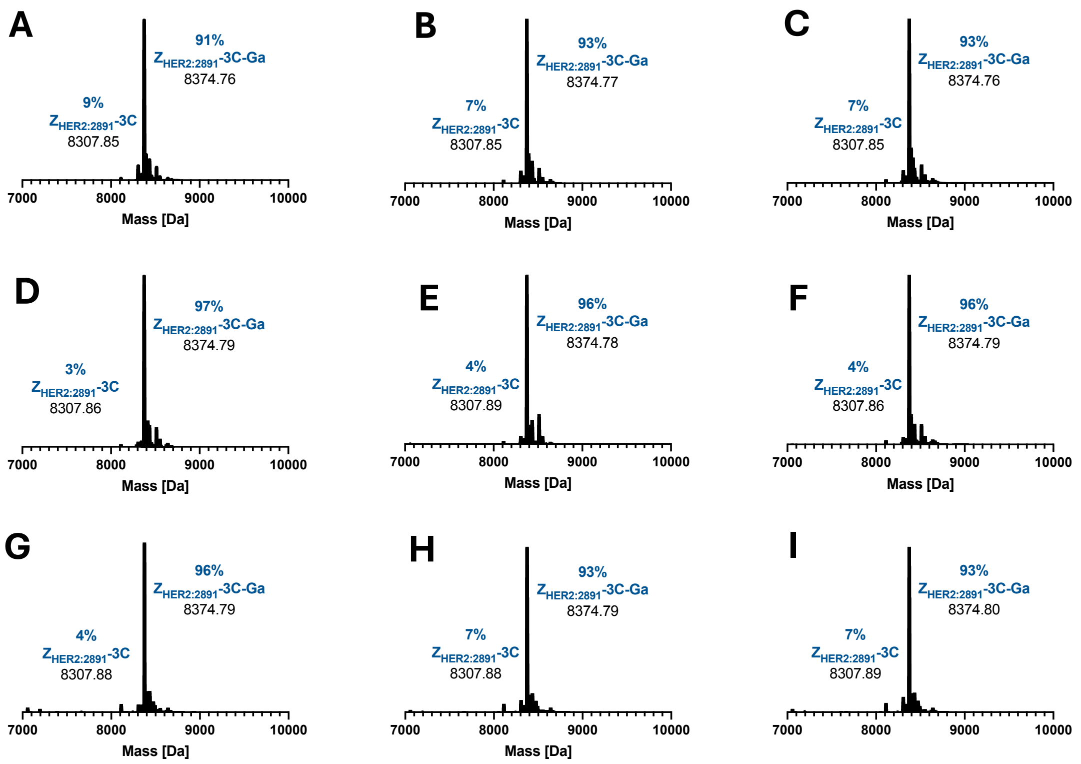


**Figure S17:** Native MS (deconvoluted) of Ga(III)-bound $Z_{HER2:2891}$-3C in triplicate stored in 100 mM ammonium acetate pH 7 at 4 °C for 0 days (A-C), 1 day (D-F) and 7 days (G-I).

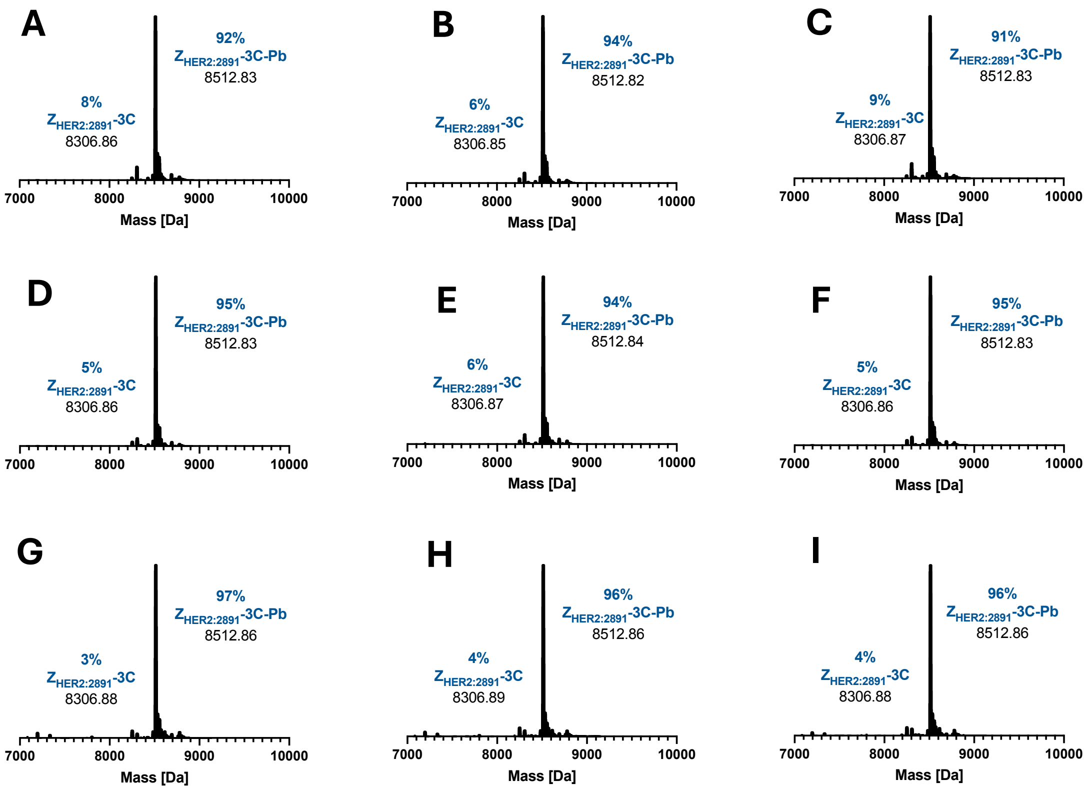


**Figure S18:** Native MS (deconvoluted) of Pb(II)-bound $Z_{HER2:2891}$-3C in triplicate stored in 100 mM ammonium acetate pH 7 at 4 °C for 0 days (A-C), 1 day (D-F) and 7 days (G-I).

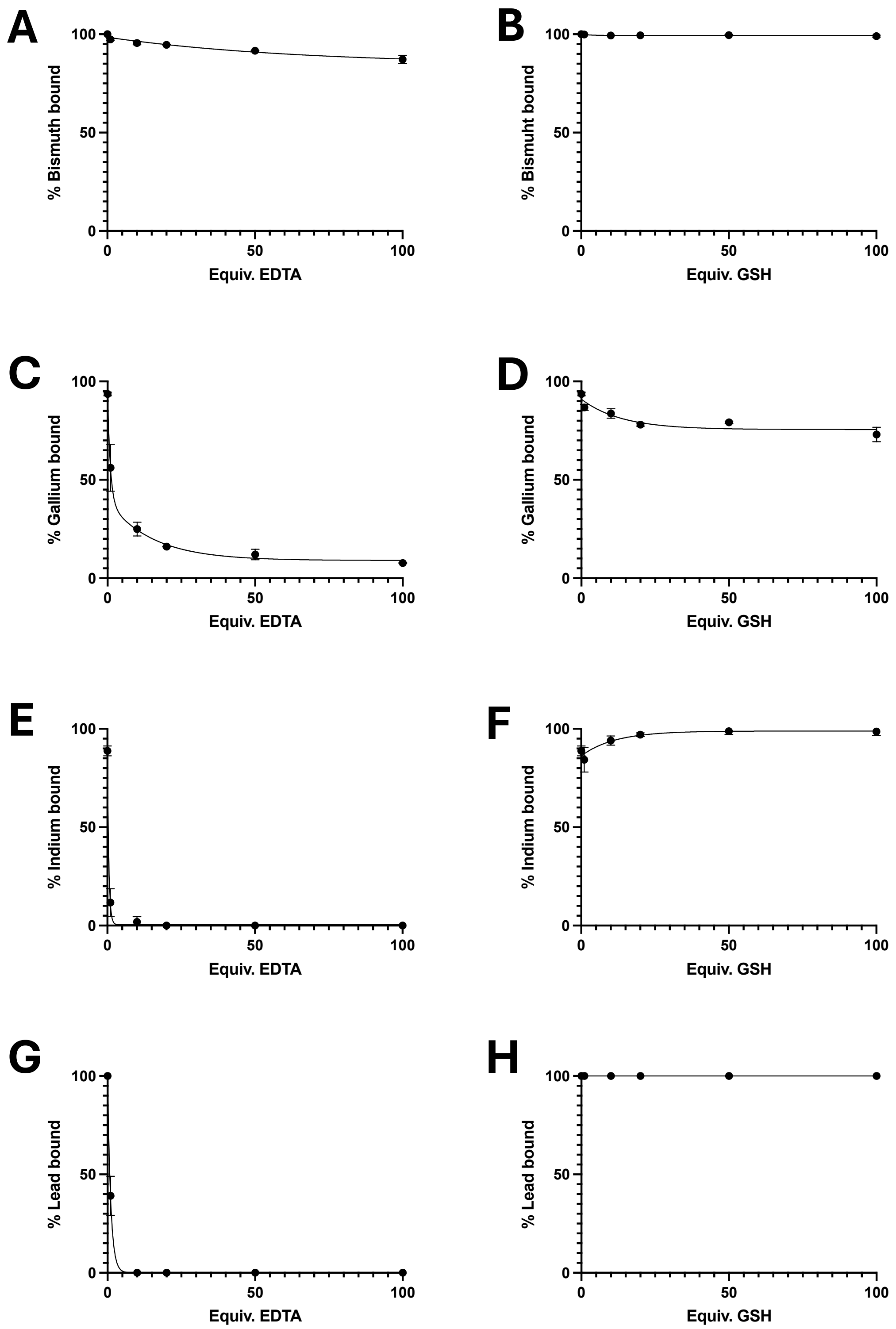


**Figure S19:** Metal retention in Bi(III)-bound $Z_{HER2:2891}$-3C in the presence of increasing equivalents of EDTA (A) and GSH (B), Ga(III)-bound $Z_{HER2:2891}$-3C in the presence of increasing equivalents of EDTA (C) and GSH (D), In(III)-bound $Z_{HER2:2891}$-3C in the presence of increasing equivalents of EDTA (E) and GSH (F), and Pb(II)-bound $Z_{HER2:2891}$-3C in the presence of increasing equivalents of EDTA (G) and GSH (H) incubated in 100 mM ammonium acetate pH 7, at 25 °C for 1 h ($n$ = 2, ± SD).

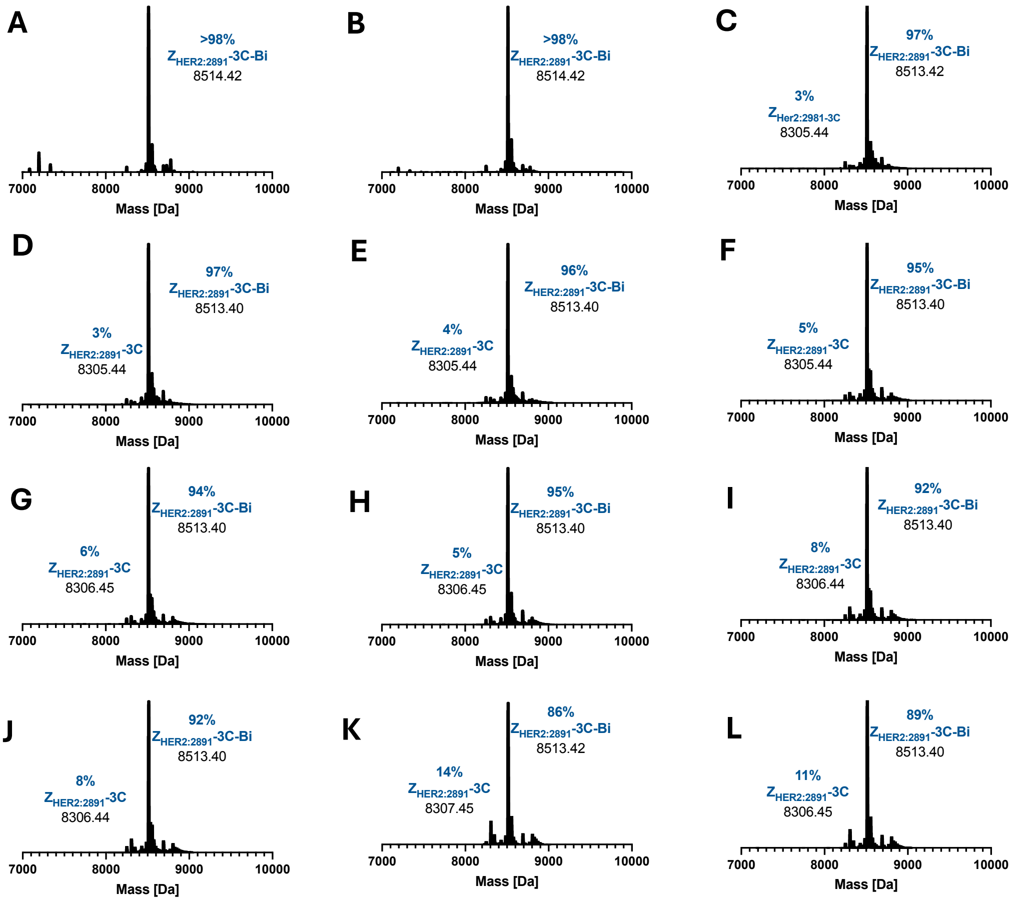


**Figure S20:** Native MS (deconvoluted) of Bi(III)-bound $Z_{HER2:2891}$-3C in duplicate after incubation for 1 h at 25 °C in the presence of 0 equiv. (A, B), 1 equiv. (C, D), 10 equiv. (E, F), 20 equiv. (G, H), 50 equiv. (I, J) or 100 equiv. (K, L) EDTA in 100 mM ammonium acetate pH 7.

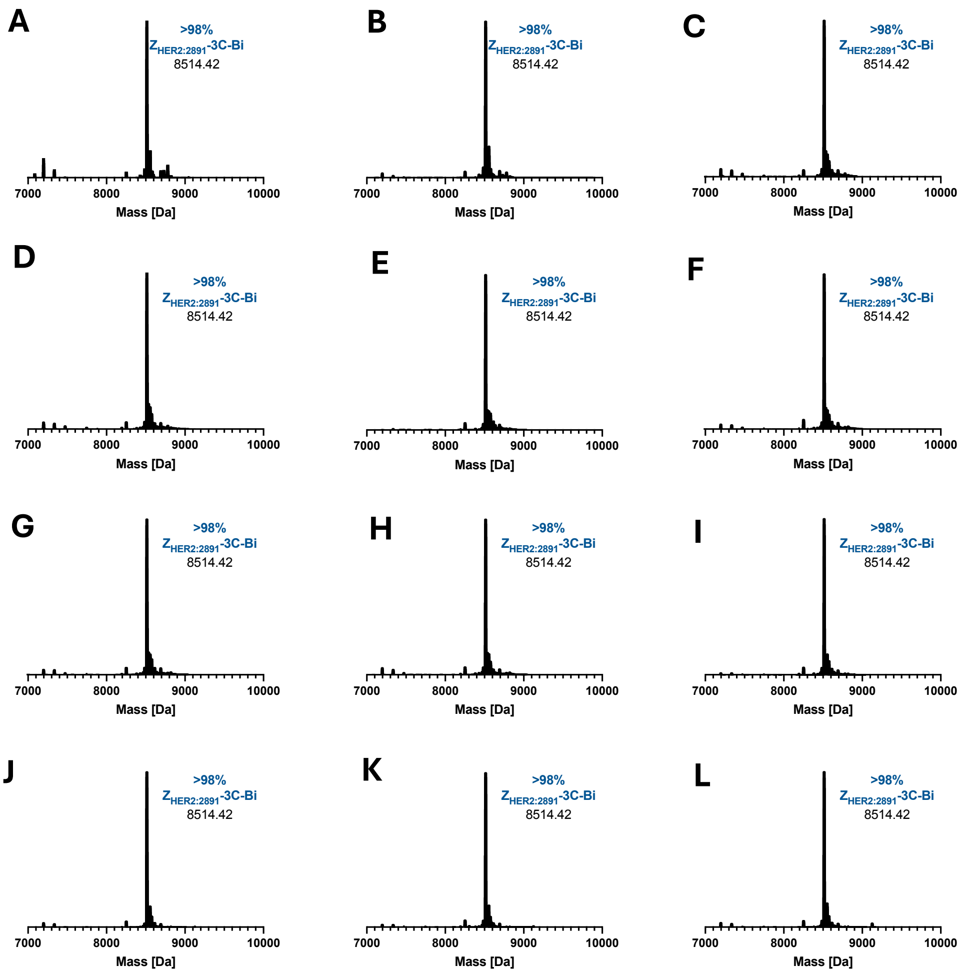


**Figure S21:** Native MS (deconvoluted) of Bi(III)-bound $Z_{HER2:2891}$-3C in duplicate after incubation for 1 h at 25 °C in the presence of 0 equiv. (A, B), 1 equiv. (C, D), 10 equiv. (E, F), 20 equiv. (G, H), 50 equiv. (I, J) or 100 equiv. (K, L) GSH in 100 mM ammonium acetate pH 7.

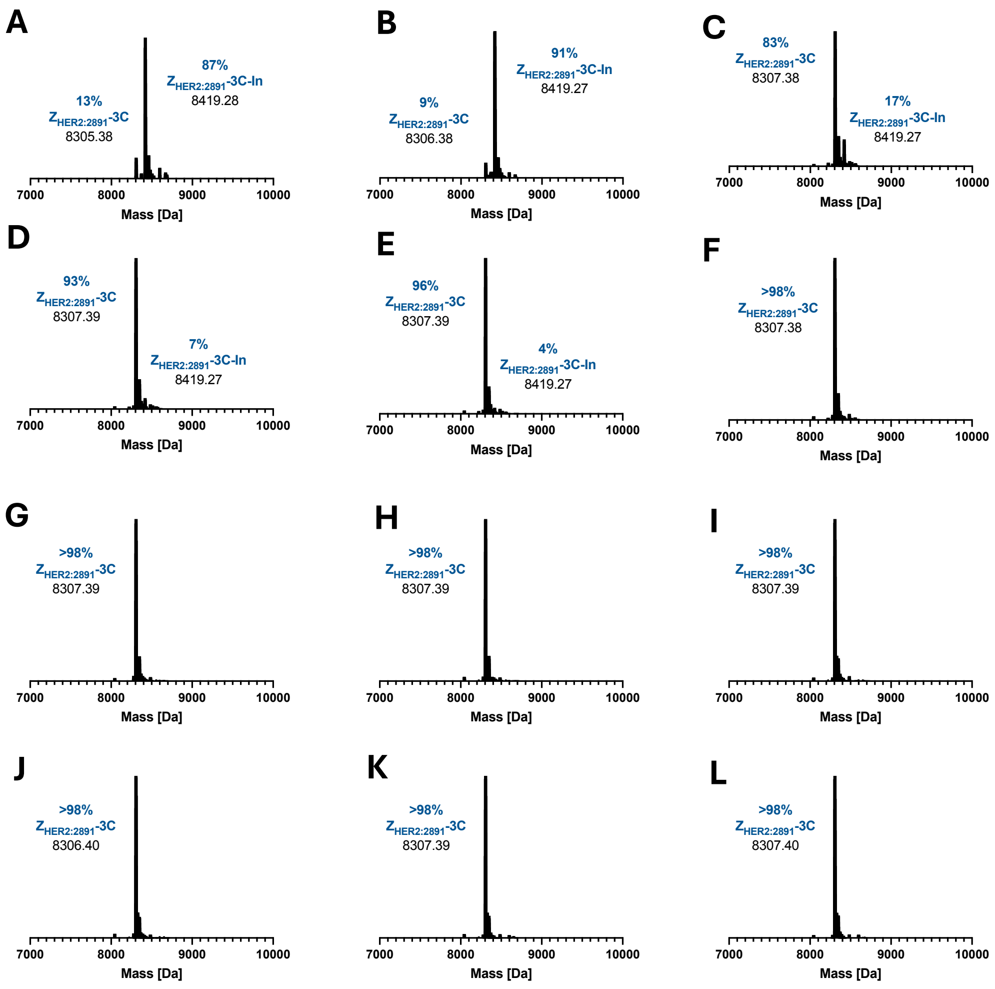


**Figure S22:** Native MS (deconvoluted) of In(III)-bound $Z_{HER2:2891}$-3C in duplicate after incubation for 1 h at 25 °C in the presence of 0 equiv. (A, B), 1 equiv. (C, D), 10 equiv. (E, F), 20 equiv. (G, H), 50 equiv. (I, J) or 100 equiv. (K, L) EDTA in 100 mM ammonium acetate pH 7.

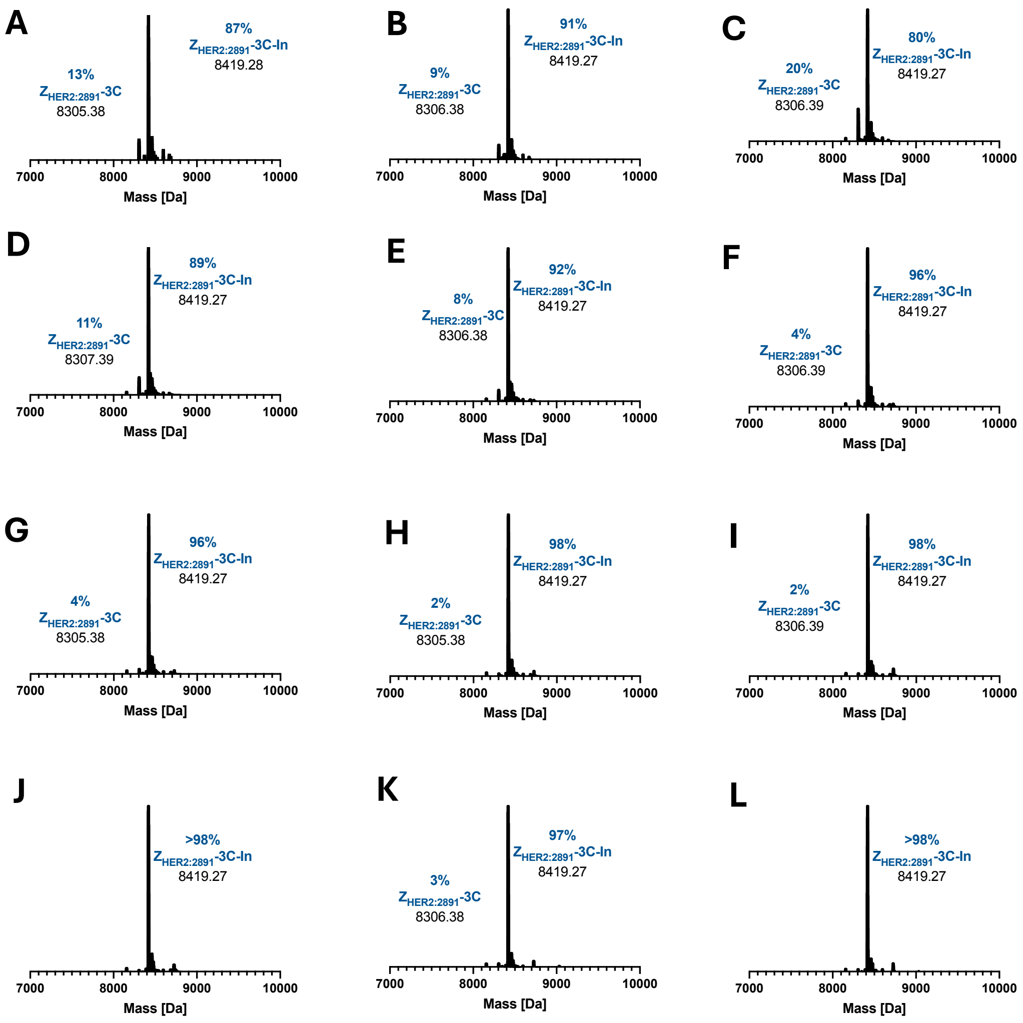


**Figure S23:** Native MS (deconvoluted) of In(III)-bound $Z_{HER2:2891}$-3C in duplicate after incubation for 1 h at 25 °C in the presence of 0 equiv. (A, B), 1 equiv. (C, D), 10 equiv. (E, F), 20 equiv. (G, H), 50 equiv. (I, J) or 100 equiv. (K, L) GSH in 100 mM ammonium acetate pH 7.

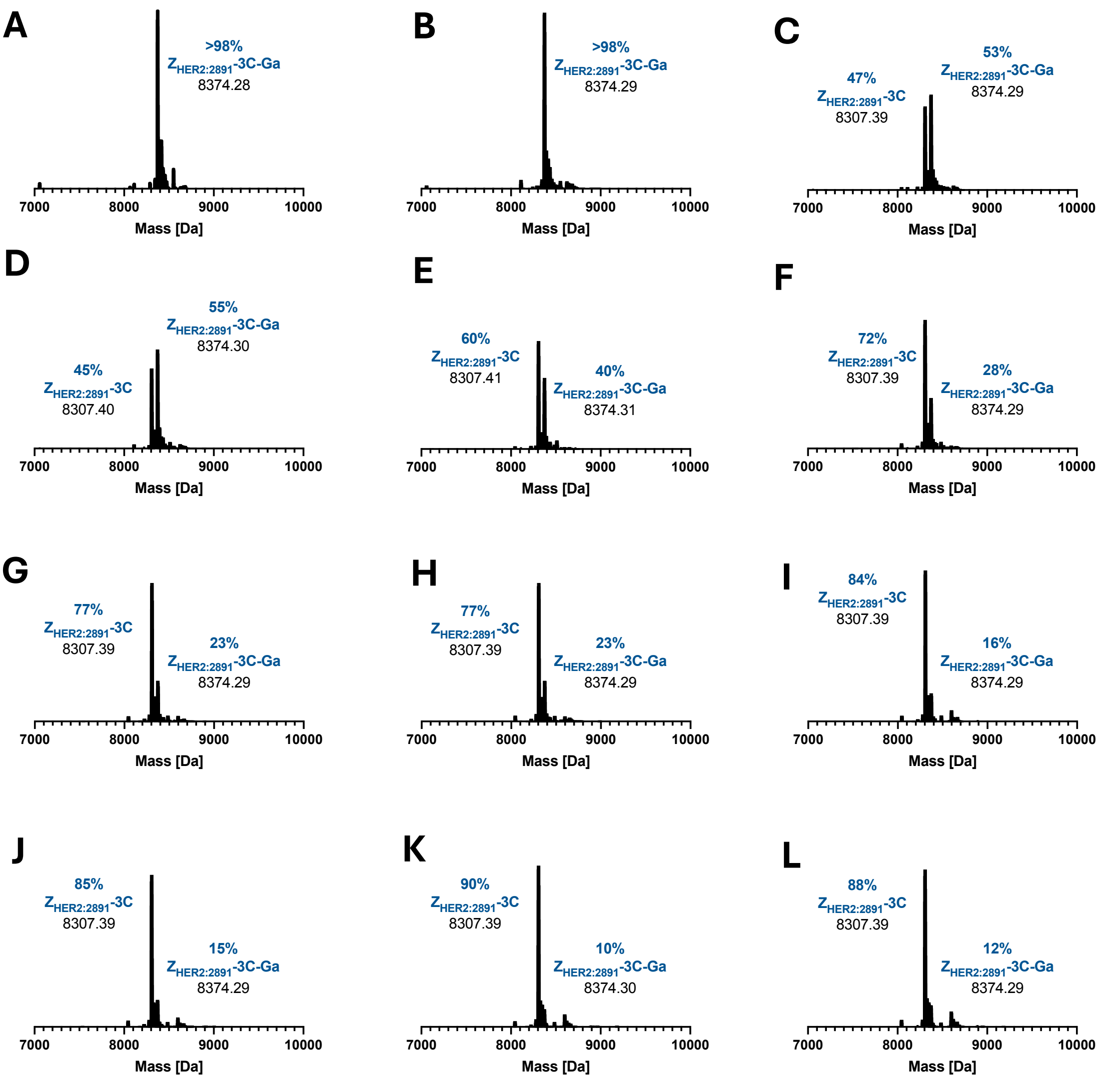


**Figure S24:** Native MS (deconvoluted) of Ga(III)-bound $Z_{HER2:2891}$-3C in duplicate after incubation for 1 h at 25 °C in the presence of 0 equiv. (A, B), 1 equiv. (C, D), 10 equiv. (E, F), 20 equiv. (G, H), 50 equiv. (I, J) or 100 equiv. (K, L) EDTA in 100 mM ammonium acetate pH 7.

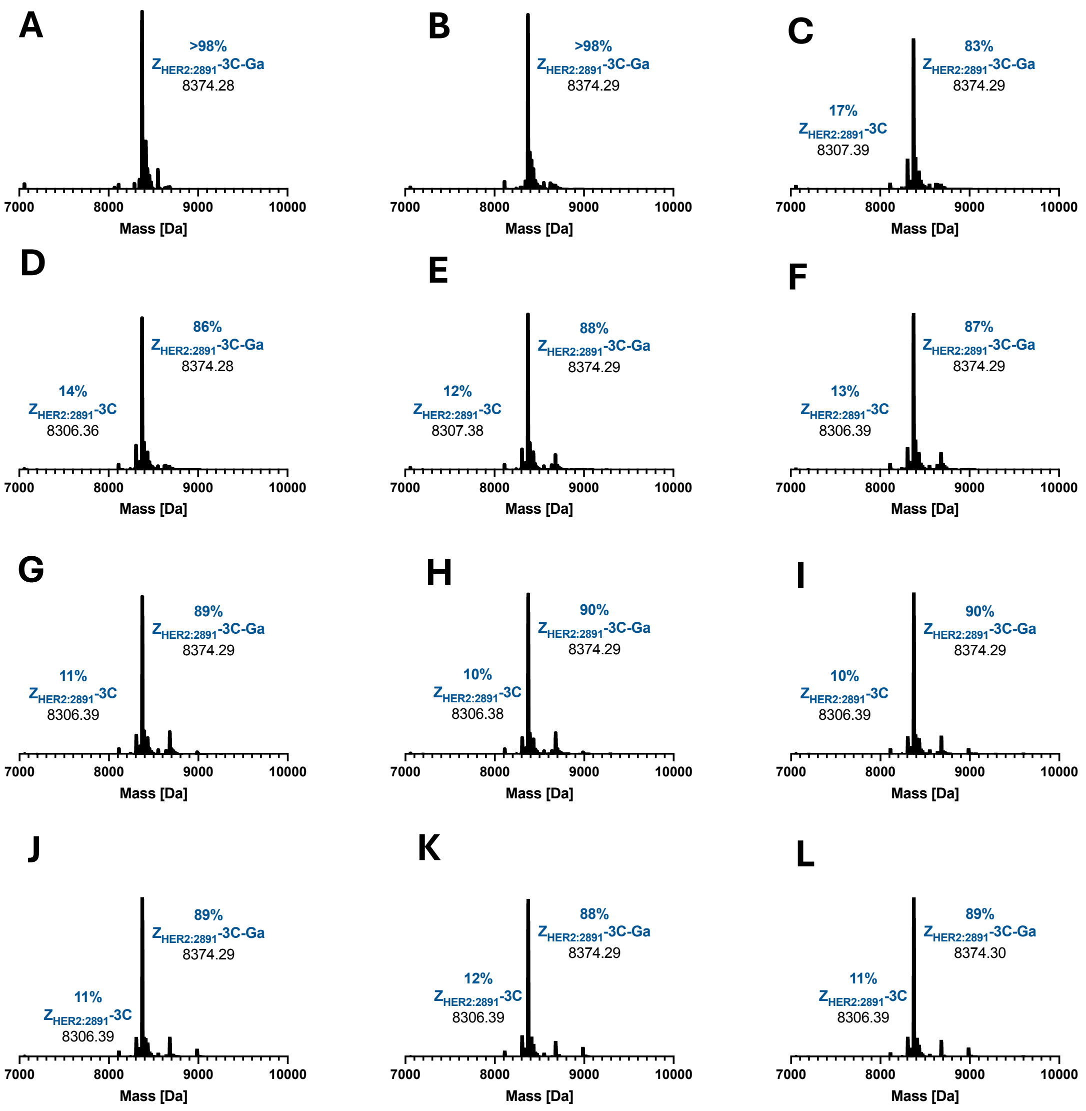


**Figure S25:** Native MS (deconvoluted) of Ga(III)-bound $Z_{HER2:2891}$-3C in duplicate after incubation for 1 h at 25 °C in the presence of 0 equiv. (A, B), 1 equiv. (C, D), 10 equiv. (E, F), 20 equiv. (G, H), 50 equiv. (I, J) or 100 equiv. (K, L) GSH in 100 mM ammonium acetate pH 7.

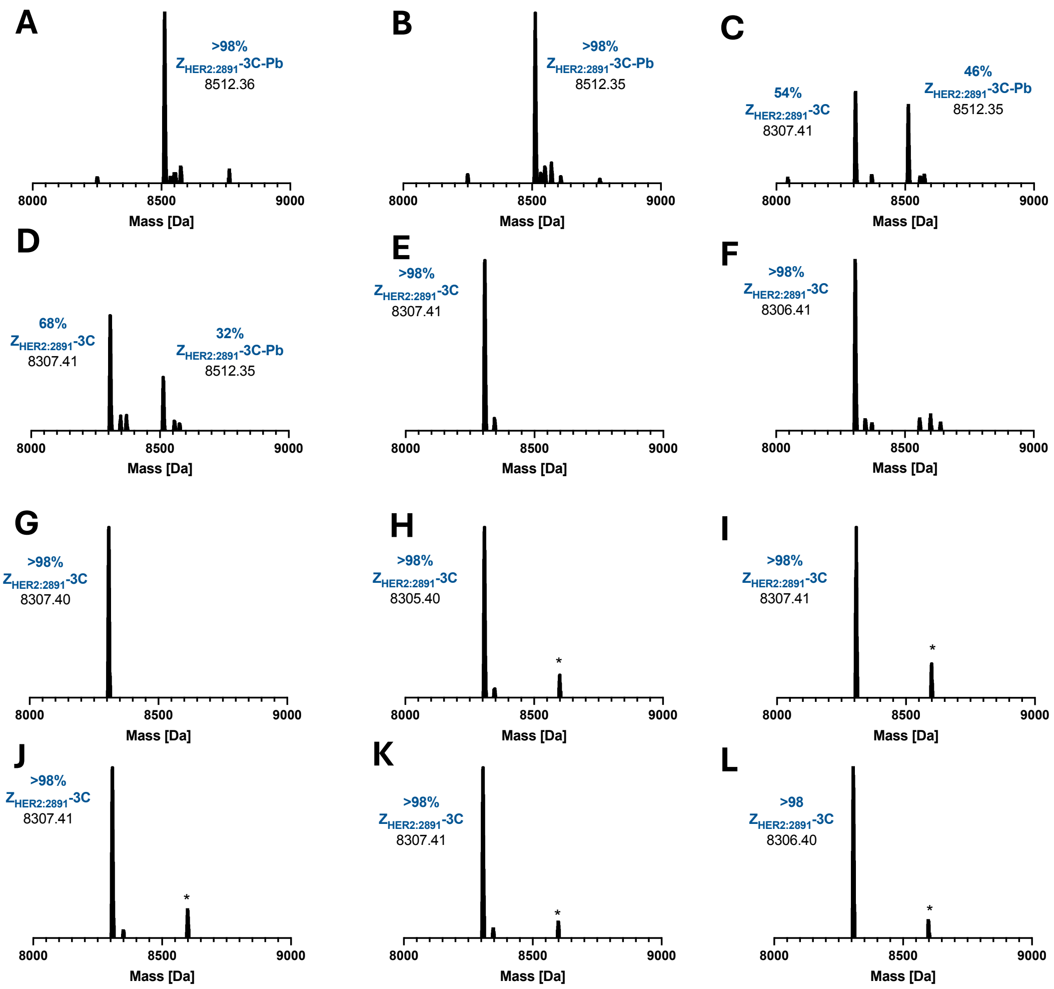


**Figure S26**: Native MS (deconvoluted) of Pb(II)-bound $Z_{HER2:2891}$-3C in duplicate after incubation for 1 h at 25 °C in the presence of 0 equiv. (A, B), 1 equiv. (C, D), 10 equiv. (E, F), 20 equiv. (G, H), 50 equiv. (I, J) or 100 equiv. (K, L) EDTA in 100 mM ammonium acetate pH 7. EDTA-Pb-$Z_{HER2:2891}$-3C adducts indicated by an asterisk (*).

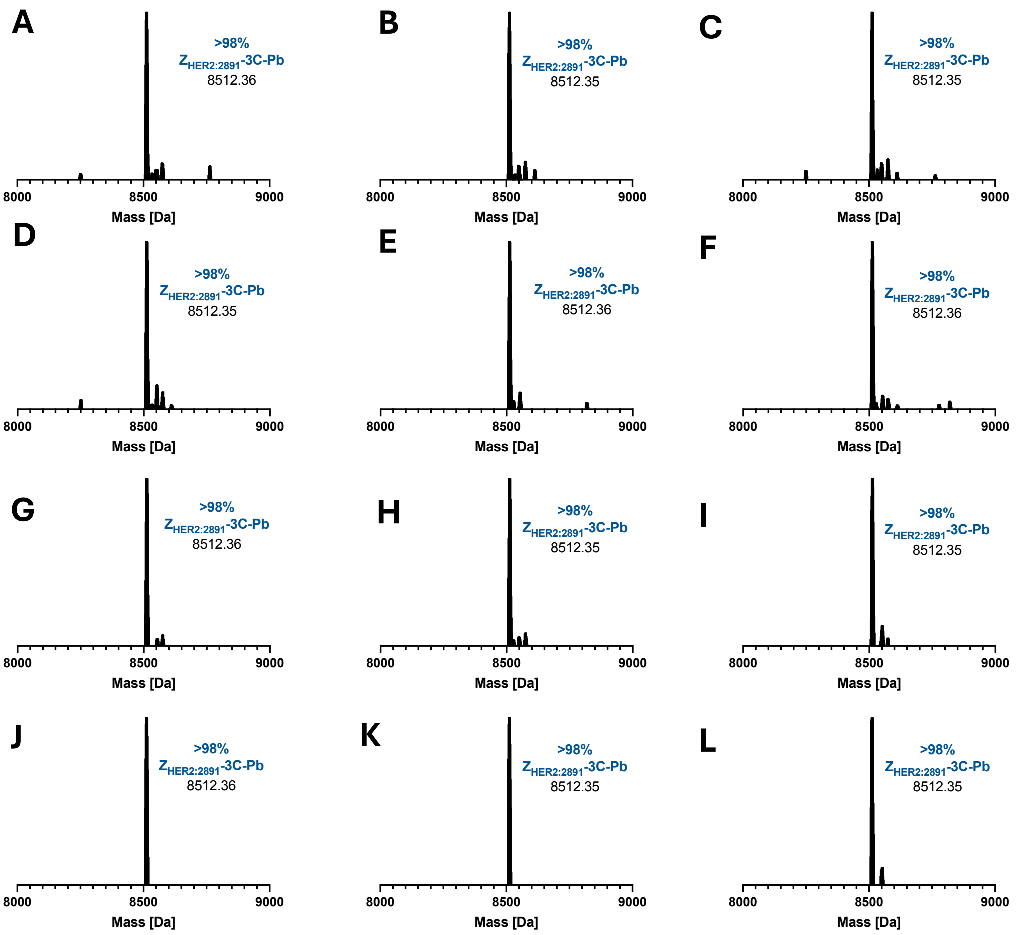


**Figure S27:** Native MS (deconvoluted) of Pb(II)-bound $Z_{HER2:2891}$-3C in duplicate after incubation for 1 h at 25 °C in the presence of 0 equiv. (A, B), 1 equiv. (C, D), 10 equiv. (E, F), 20 equiv. (G, H), 50 equiv. (I, J) or 100 equiv. (K, L) GSH in 100 mM ammonium acetate pH 7.

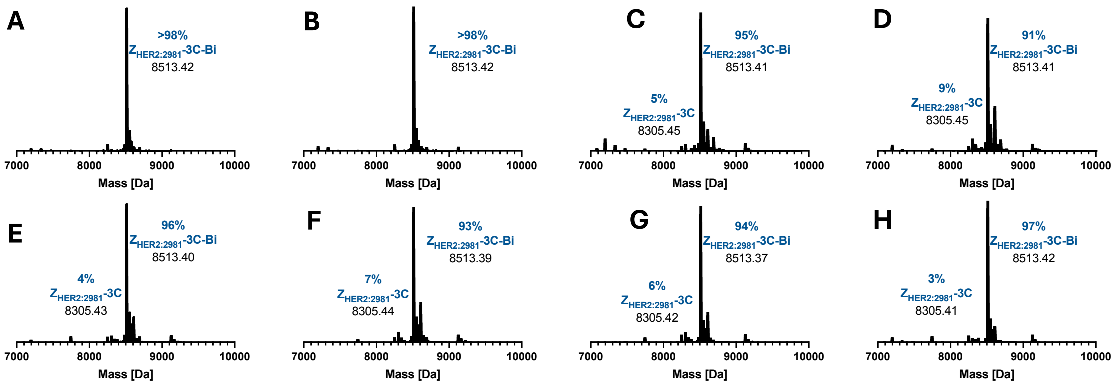


**Figure S28:** Native MS (deconvoluted) of Bi(III)-bound $Z_{HER2:2891}$-3C in duplicate after incubation for 1 h at 25 °C in the presence of 100 eq GSH in 100 mM ammonium acetate pH 7 then subsequent storage at 4 °C for 1 day (A, B), 5 days (C, D), 8 days (E,F) and 14 days (G, H).

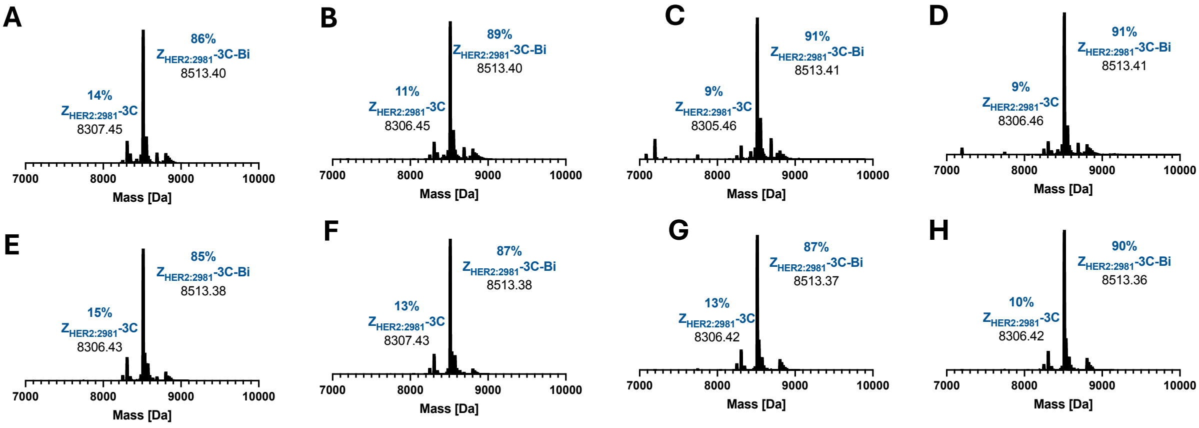


**Figure S29:** Native MS (deconvoluted) of Bi(III)-bound $Z_{HER2:2891}$-3C in duplicate after incubation for 1 h at 25 °C in the presence of 100 eq EDTA in 100 mM ammonium acetate pH 7 then subsequent storage at 4 °C for 1 day (A, B), 5 days (C, D), 8 days (E,F) and 14 days (G, H).

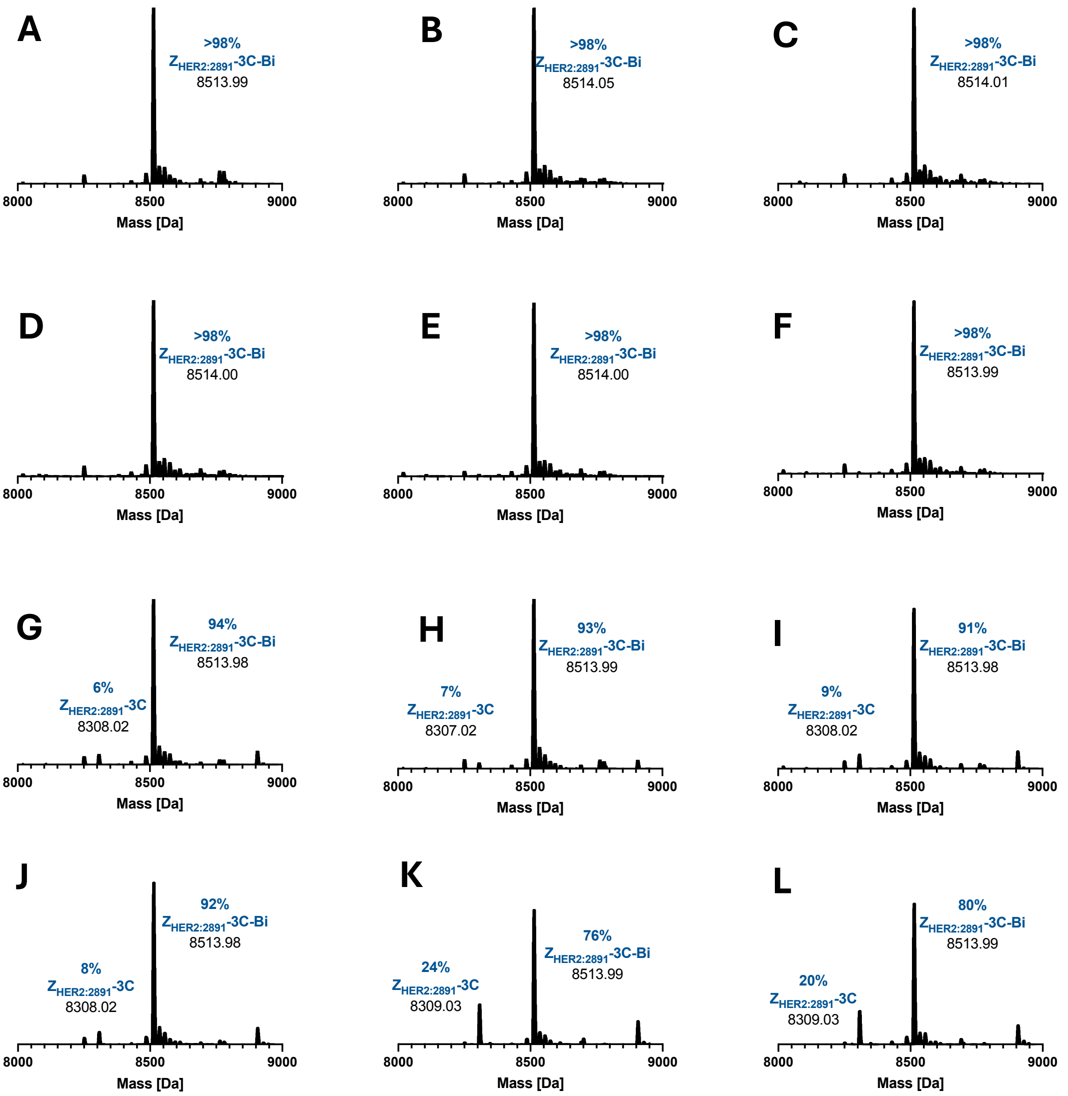


**Figure S30:** Native MS (deconvoluted) of Bi(III)-bound $Z_{HER2:2891}$-3C in duplicate after incubation for 1 h at 25 °C in the presence of 0 equiv. (A, B), 1 equiv. (C, D), 10 equiv. (E, F), 20 eq (G, H), 50 equiv. (I, J) or 100 equiv. (K, L) DTPA in 100 mM ammonium acetate pH 7.

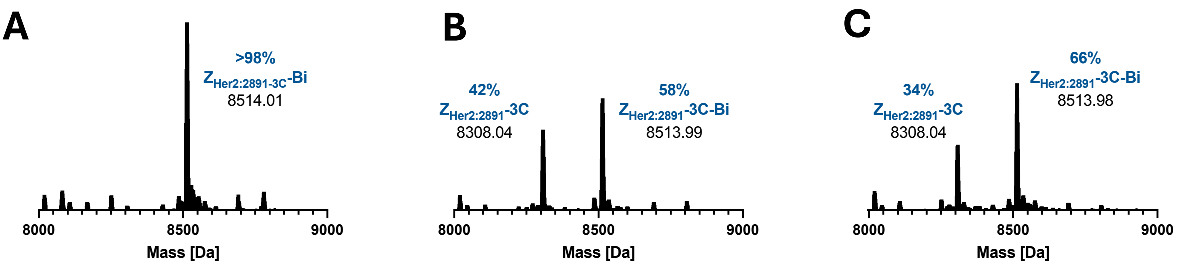


**Figure S31:** Native MS (deconvoluted) of Bi(III)-bound $Z_{HER2:2891}$-3C after incubation for 10 minutes at 100 °C in the presence of 0 equiv. (A) or 100 equiv. (B, C) EDTA in 100 mM ammonium acetate pH 7.

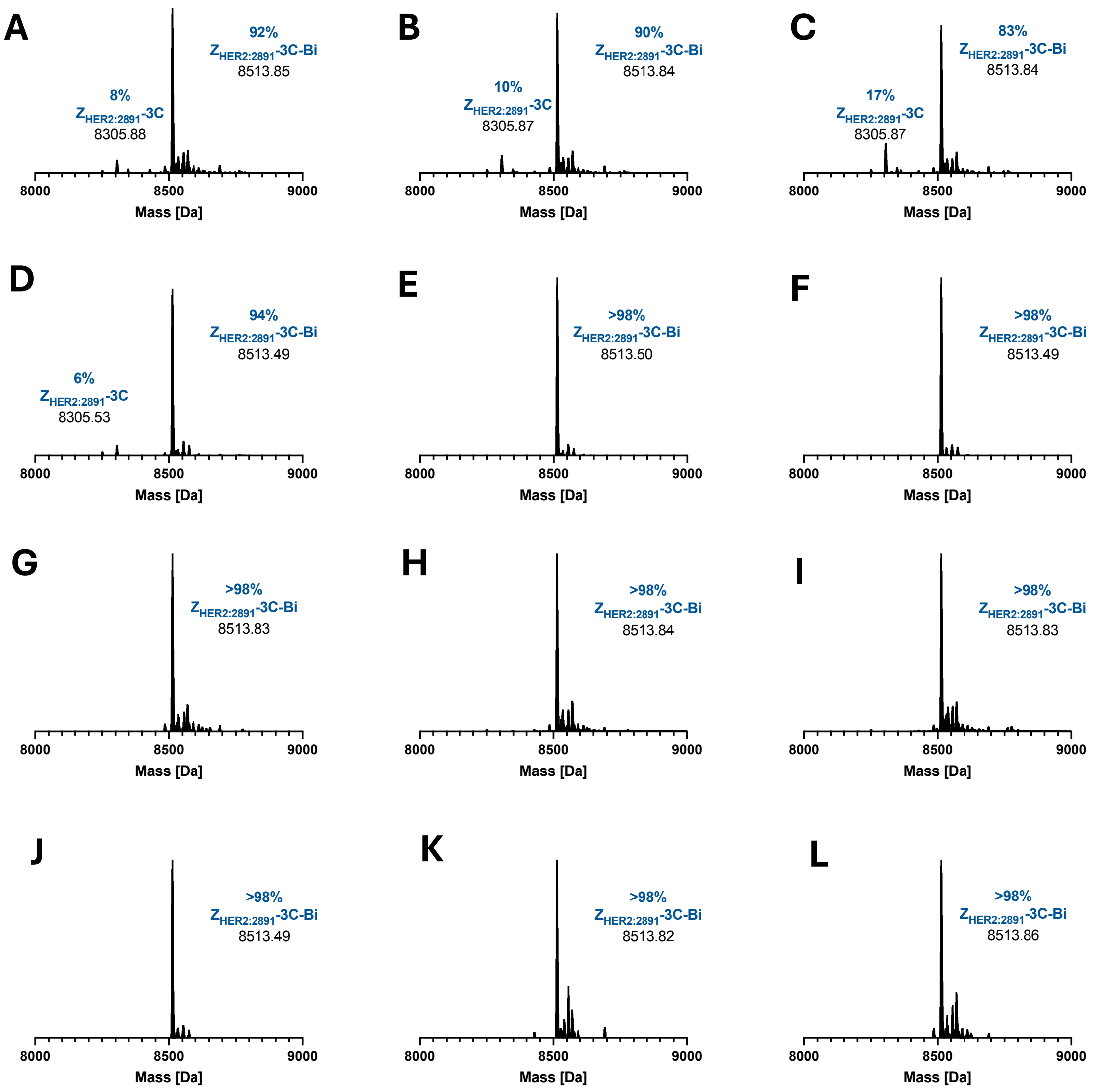


**Figure S32**: Native MS (deconvoluted) of the optimisation of Bi(III) uptake into $Z_{HER2:2891}$-3C after incubation in 50 mM TCEP, 150 mM NaCl, 10 mM Tris at pH 2, 25 °C for 15 (A), 30 (B) and 60 (C) minutes, or pH 2, 67 °C for 15 (D), 30 (E) and 60 (F) minutes, or pH 7.5, 25 °C for 15 (G), 30 (H) and 60 (I) minutes, or pH 7.5, 67 °C for 15 (J), 30 (K) and 60 (L) minutes, prior to metal addition and buffer exchange into 100 mM ammonium acetate, pH 7.

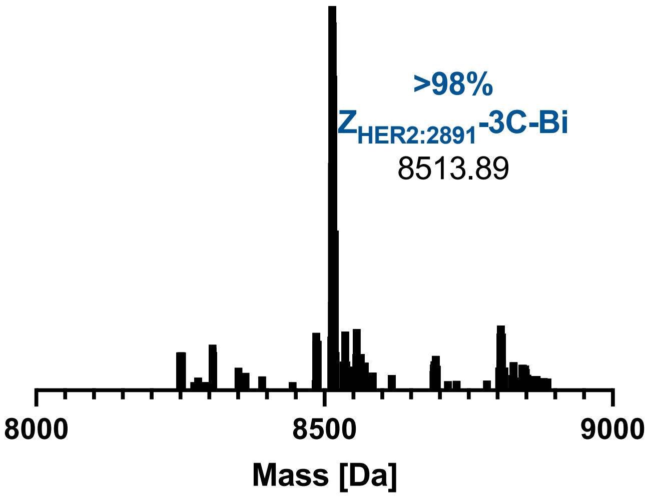


**Figure S33:** Native MS (deconvoluted) of Bi(III)-bound $Z_{HER2:2891}$-3C after incubation for 1 h at 25 °C in the presence of 100 equiv. EDTA in 100 mM ammonium acetate pH 7. Collected with an alternative MS source (Synapt alternative native MS method as described above).

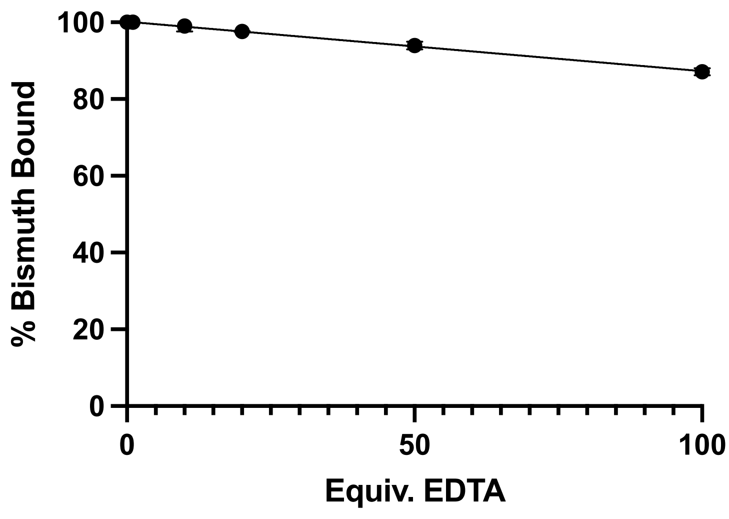


**Figure S34:** Metal retention in Bi(III)-bound $Z_{TNF\text{-}alpha1}$-3C in the presence of increasing equivalents of EDTA, incubated in 100 mM ammonium acetate pH 7, at 25 °C for 1 h ($n = 2$, ± SD).

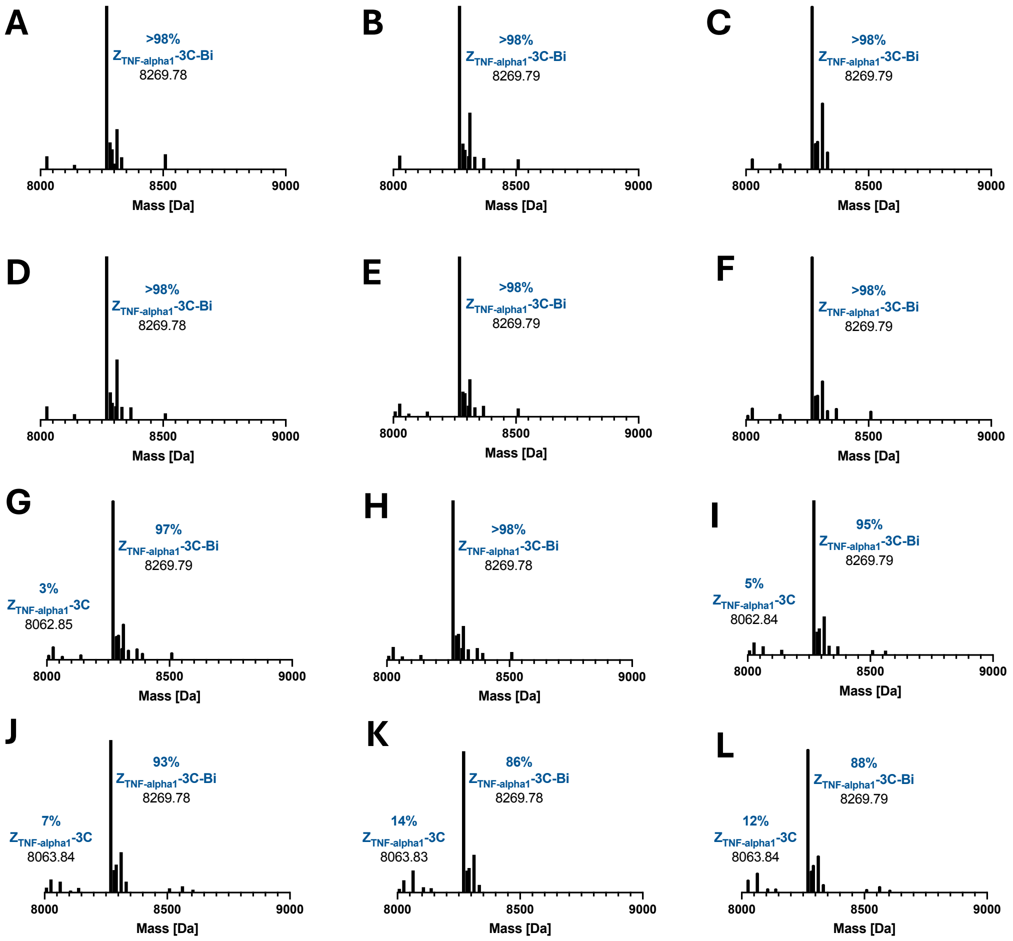


**Figure S35:** Native MS (deconvoluted) of Bi(III)-bound $Z_{TNF\text{-}alpha1}$-3C in duplicate after incubation for 1 h at 25 °C in the presence of 0 equiv. (A, B), 1 equiv. (C, D), 10 equiv. (E, F), 20 equiv. (G, H), 50 equiv. (I, J) or 100 equiv. (K, L) EDTA in 100 mM ammonium acetate pH 7.

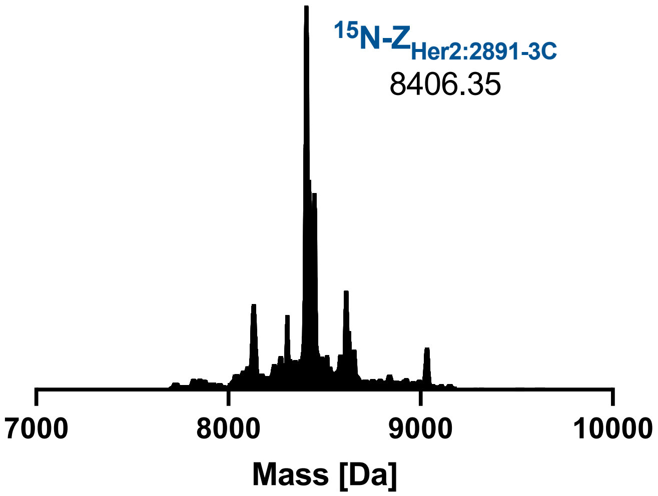


**Figure S36:** Intact MS of $^{15}$N labelled $Z_{HER2:2891}$-3C in in 20 mM MES pH 7.5, 150 mM NaCl.

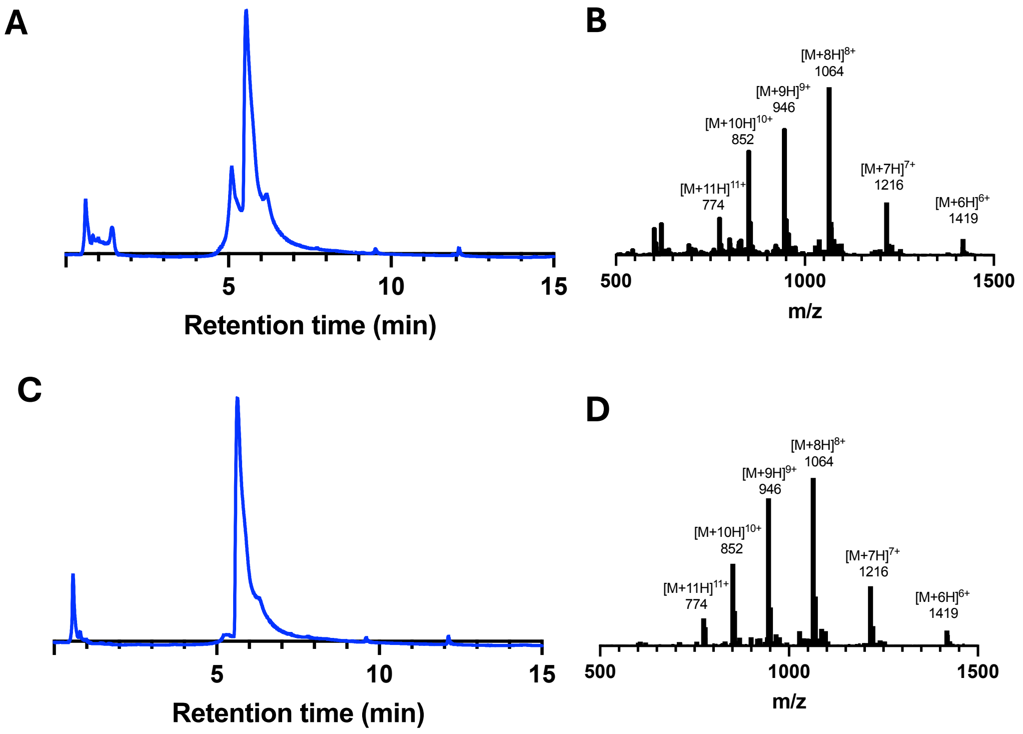


**Figure S37:** Analytical LCMS (5-95% ACN:$H_2O$ with 0.1% formic acid (*v/v*) over 15 minutes) of lyophilised $Z_{HER2:2891}$-3C-Bi after redissolving and metal uptake via the optimised metal uptake protocol. (A, C) UV traces at 280 nm where the metal bound affibody elutes at ~6 minutes. (B, D) mass spectra corresponding to the affibody peak in the LC.

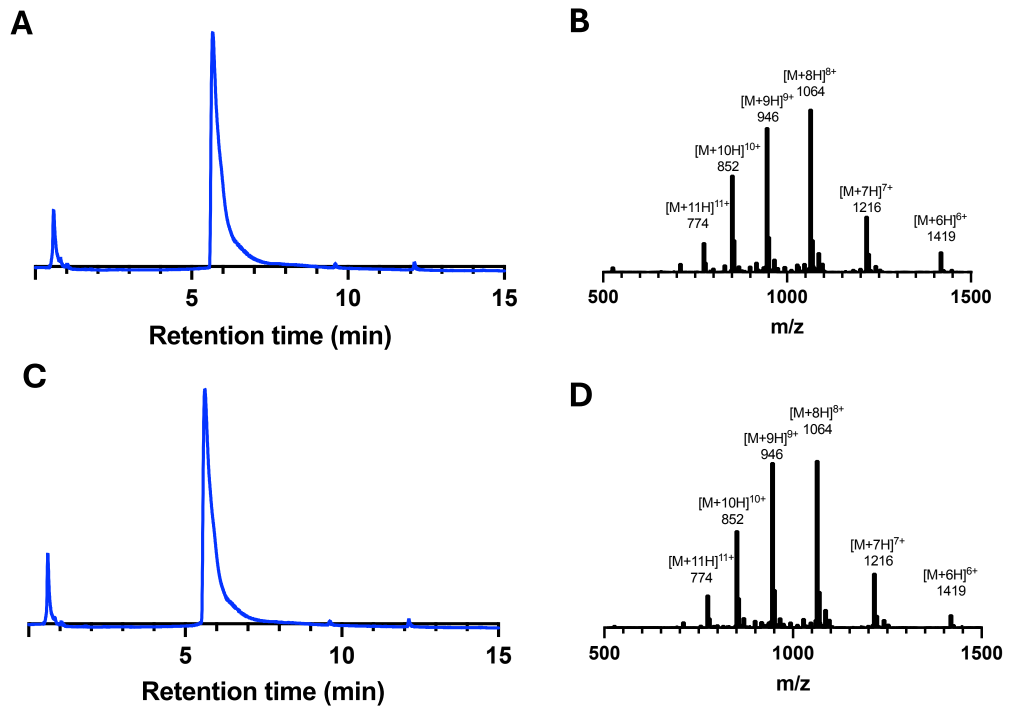


**Figure S38:** Analytical LCMS (5-95% ACN:$H_2O$ with 0.1% formic acid (*v/v*) over 15 minutes) of $Z_{HER2:2891}$-3C-Bi after metal uptake via the optimised metal uptake protocol. (A, C) UV traces at 280 nm where the metal bound affibody elutes at ~6 minutes. (B, D) mass spectra corresponding to the affibody peak in the LC.

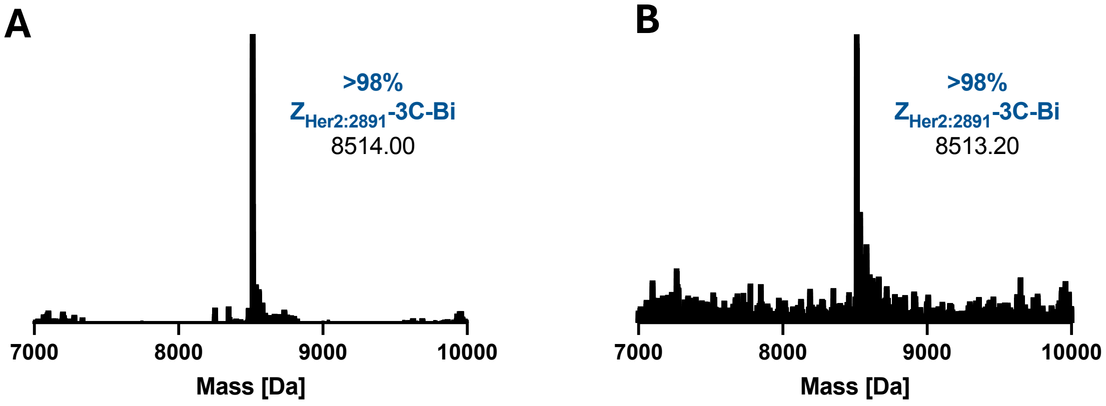


**Figure S39:** Native MS (deconvoluted) of lyophilised $Z_{HER2:2891}$-3C after redissolving and metal uptake via the optimised metal uptake protocol collected with the orthogonal native MS method.

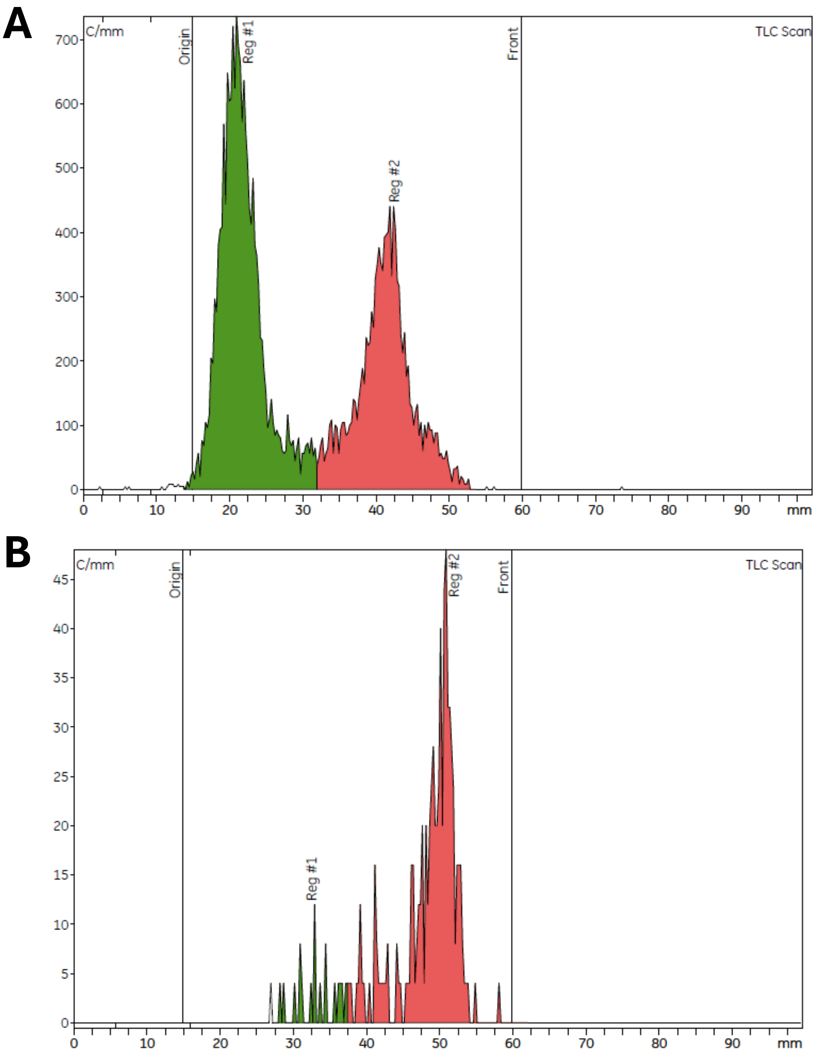


**Figure S40:** TLC chromatograms of the crude reaction mixture of $Z_{Her2:2891}$-3C and $^{225}AcCl_3$. (A) t = 0 showing $Z_{Her2:2891}$-3C-$^{213}$Bi at the start (origin) of the chromatogram and free radionuclides at the front. (B) t = 5 days showing free radionuclides at the front of the chromatogram.

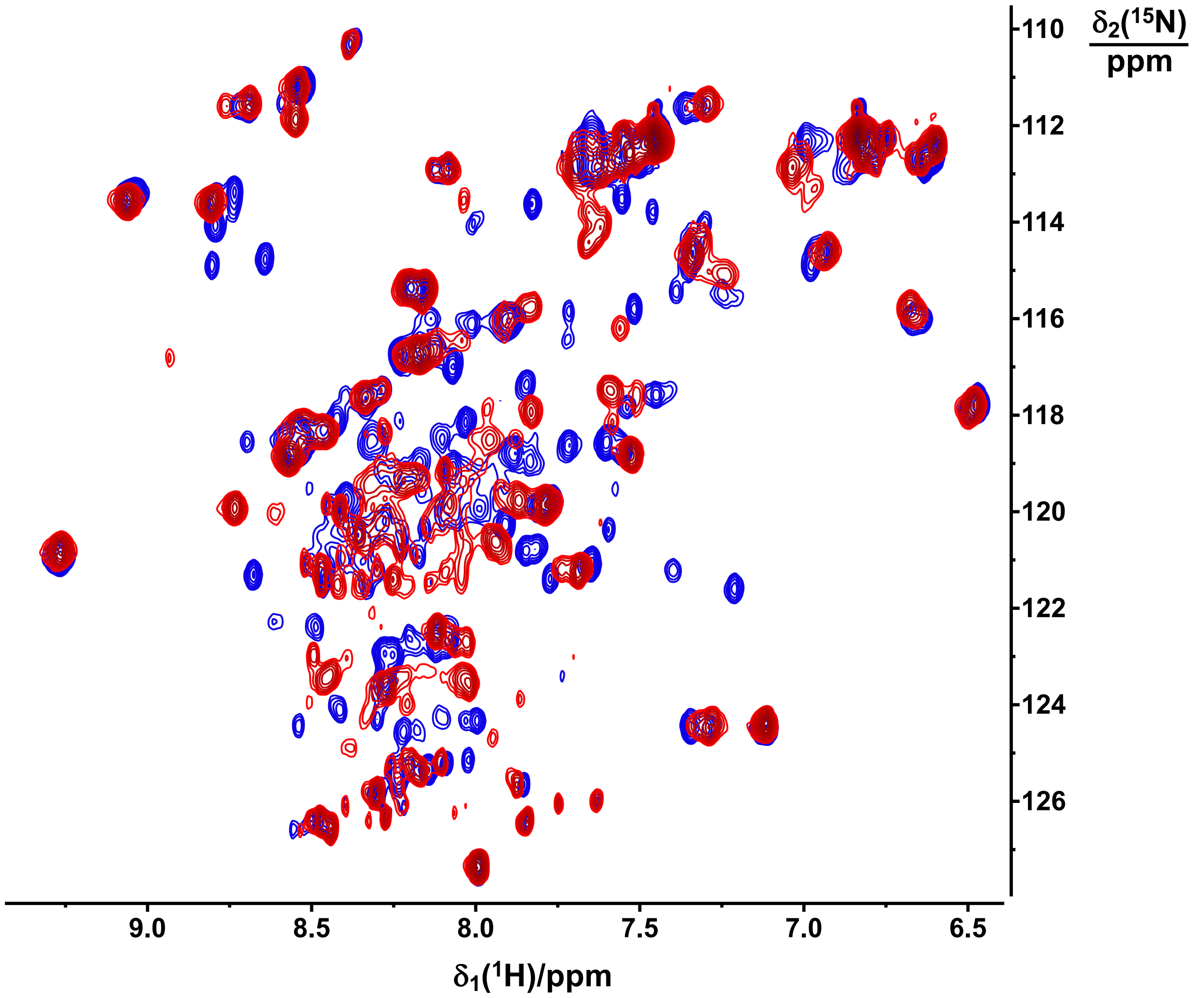


**Figure S41:** Superimposed 800 MHz [$^{15}$N,$^{1}$H]-HSQC NMR spectra (Watergate by shaped pulses water suppression) of 400 µM solutions of reduced $^{15}$N-Z$_{HER:2891}$-3C apo (blue) and $^{15}$N-Z$_{HER:2891}$-3C in presence 1 eq. Bi(III) (red) in 20 mM MES pH 7.5, 150 mM NaCl, 20 mM TCEP, 10% $D_2O$.

[4] John Jumper, Richard Evans, Alexander Pritzel, Tim Green, Michael Figurnov, Olaf Ronneberger, Kathryn Tunyasuvunakool, Russ Bates, Augustin Žídek, Anna Potapenko, Alex Bridgland, Clemens Meyer, Simon A. A. Kohl, Andrew J. Ballard, Andrew Cowie, Bernardino Romera-Paredes, Stanislav Nikolov, Rishub Jain, Jonas Adler, Trevor Back, Stig Petersen, David Reiman, Ellen Clancy, Michal Zielinski, Martin Steinegger, Michalina Pacholska, Tamas Berghammer, Sebastian Bodenstein, David Silver, Oriol Vinyals, Andrew W. Senior, Koray Kavukcuoglu, Pushmeet Kohli, D. Hassabis, *Nature* **2021**, *596*, 583–589.

[5] Y. J. Wang, L. Xu, *J. Inorg. Biochem.* **2008**, *102*, 988–991.

[6] E. C. Meng, T. D. Goddard, E. F. Pettersen, G. S. Couch, Z. J. Pearson, J. H. Morris, T. E. Ferrin, *Protein Sci.* **2023**, *32*, e4792.
